# Bayesian model selection reveals biological origins of zero inflation in single-cell transcriptomics

**DOI:** 10.1101/2020.03.03.974808

**Authors:** Kwangbom Choi, Yang Chen, Daniel A. Skelly, Gary A. Churchill

## Abstract

Single-cell RNA sequencing is a powerful tool for characterizing cellular heterogeneity in gene expression. However, high variability and a large number of zero counts present challenges for analysis and interpretation. There is substantial controversy over the origins and proper treatment of zeros and no consensus on whether zero-inflated count distributions are necessary or even useful. While some studies assume the existence of zero inflation due to technical artifacts and attempt to impute the missing information, other recent studies of argue that there is no zero inflation in scRNA-Seq data. We apply a Bayesian model selection approach to unambiguously demonstrate zero inflation in multiple biologically realistic scRNA-Seq datasets. We show that the primary causes of zero inflation are not technical but rather biological in nature. We also demonstrate that parameter estimates from the zero-inflated negative binomial distribution are an unreliable indicator of zero inflation. Despite the existence of zero inflation of scRNA-Seq counts, we recommend the generalized linear model with negative binomial count distribution (not zero-inflated) as a suitable reference model for scRNA-Seq analysis.

---

Single-cell RNA Sequencing (scRNA-Seq) is a powerful tool for studying the dynamics of gene expression and for characterizing heterogeneity in complex mixtures of cells. Technical advances, including unique molecular identifiers (UMIs) [Islam et al., 2014], combinatorial barcoding [Cao et al., 2017,Rosenberg et al., 2018], and physical containment of cells in droplets [Macosko et al., 2015,Klein et al., 2015] have enabled profiling of ever larger numbers of cells with fewer RNA molecules sequenced per cell. The sparseness of single cell data presents challenges for analysis and interpretation. In particular, the high proportion of zero counts (zero inflation) that is observed for many genes has become a major focus of discussion and debate. Zeros have variously been attributed to technical artifacts [Kharchenko et al., 2014,Hicks et al., 2017] or to statistical sampling [Townes et al., 2019,Svensson, 2020]. Less attention has been given to the biological factors that might contribute to zero inflation including the role of cellular heterogeneity. Substantial controversy has ensued over approaches for mitigating potential bias rooted in zero inflation. In particular, numerous approaches have been proposed to replace observed zeros in count data with imputed non-zero values based on the assumption that zeros are due to technical artifacts [Gong et al., 2018,Li and Li, 2018]. On the other hand, several recent studies use negative control data to demonstrate that the occurrence of zeros is consistent with expectations from statistical sampling [Townes et al., 2019,Svensson, 2020] and implicate against imputation. In order to resolve these conflicting views, a principled examination of zeros in biologically realistic scRNA-Seq data is needed.

Statistical distributions describe the expected frequency of counts, including zeros, under specific assumptions about the data generating process. The most widely used distribution for count data is the Poisson (P) distribution. In the context of scRNA-Seq data, the Poisson distribution follows from the assumptions that each cell has an equal proportion of a given mRNA species and that the observed counts reflect independent statistical sampling across cells. Count data are often more variable than predicted by the Poisson distribution. This has led to the adoption of count distributions such as the negative binomial (NB) that include an overdispersion parameter, *r*, to account for this extra variation. Count distributions can be extended to include a zero-inflated component that generates zeros at random (with probability *π*_0_) regardless of the actual amount of mRNA present in a cell. One can imagine that some of the counts that could have been non-zero are masked and replaced by zeros. Zero-inflated Poisson (ZIP) and zero-inflated negative binomial (ZINB) distributions are commonly used in this setting.

In addition to the count distribution, a statistical model of scRNA-Seq data should be able to incorporate explanatory variables to account for sex, cell type, or treatment effects that are the focus of the experimental investigation as well as batch effects. Generalized linear models (GLMs) for count data are well-established statistical tools that provide sophisticated modeling and inference capabilities using off-the-shelf software [Zeileis et al., 2008,Bürkner, 2017,Goodrich et al., 2019]. GLMs are directly applicable to count data and do not require preprocessing such as scaling, normalization, or log-transformation with pseudo-counts. A GLM developed for scRNA-Seq data should include an adjustment (offset) that accounts for cell-to-cell variation in the depth of sequencing. The offset effectively normalizes the data with respect to variation in total UMI count per cell without directly altering the data. In particular, the zero counts remain as zeros. The effect of including an offset is to convert the scale of the GLM from a model of expected counts to a model of expected rates of expression (*µ*) that is comparable across cells with different total UMI counts.

Using the GLM framework, we apply a Bayesian model selection criterion [Vehtari et al., 2017] to scRNA-Seq data to identify the statistical distributions that best fit the data for each gene, including zero and non-zero values. This is a more comprehensive evaluation of zero inflation than previous studies that have relied solely on the comparison of the observed versus expected proportion of zeros after model fitting [Townes et al., 2019,Svensson, 2020]. We consider the implications of statistical sampling, technical dropout, cell heterogeneity, and key biological variables, and compare these to the observed data to better understand the statistical properties of zero inflation and to evaluate the impact of different modeling choices on inference.

Typical droplet scRNA-Seq experiments utilize UMI counts to quantify gene expression in single cells [Islam et al., 2014]. We center our analysis around data from a droplet scRNA-Seq experiment that shares characteristics common to many recent scRNA-Seq experiments. To generate these data, Skelly et al. [Skelly et al., 2018] used 10X Chromium technology to profile cells from the non-myocyte fraction of female and male mouse hearts. Cardiac non-myocytes, which predominantly include leukocytes, vascular cells, and stromal cells, exhibit considerable transcriptional and cellular diversity. Like other recent scRNA-Seq experiments these data include transcriptional profiles of thousands of cells (10,519) sequenced at a relatively low per-cell depth (median 4,270 UMIs) and display a high total fraction of zeros (>93%). The findings we report are general and are apparent in other scRNA-Seq datasets. In the Supplemental Materials we report the analysis of experimental data obtained from two additional biologically heterogeneous datasets, namely mouse kidney [Park et al., 2018] and human peripheral blood mononuclear cells [10X Genomics, 2018].

## Results

### Why are there so many zeros?

The most important factor that determines the number of zeros in scRNA-Seq data is the sequencing depth (total UMI count) per cell. In the heart data, this ranges from 746 to 17,302 UMIs per cell after filtering to 5,515 genes in order to remove genes with non-zero UMI counts in less than 10% of cells (Figure1a, Supplemental FigureS1a and S1b). Clearly, if the total UMI count in a cell is less than the number of genes, some genes will have zero counts. Sequencing depth explains 95% of variation in the number of zeros per cell (*R*^2^ = 0.945, *p <* 2.2*e^−^*^16^). We account for this variation by including an offset term in the GLM, which we have scaled to represent rates of expression in units of UMI/10K.

The second most important factor in determining the number of zeros is the per-gene average rate of expression. In the heart data the rate of expression varies from 0.23 to 97.4 UMI/10K across 5,515 genes (Figure1b; Supplemental FigureS1c). In general, genes with lower rates of expression will have a higher frequency of zeros. We compared the expected number zeros for each gene, assuming a Poisson model with matching gene-specific rates of expression and cellspecific offsets, to the observed numbers of zeros in the heart data (Supplemental FigureS1d). We see that many genes have “extra” zeros, in some cases with thousands of zeros over expectation. Presumably, these genes have count distributions that are overdispersed (NB), zero-inflated (ZIP), or both (ZINB).

**Figure 1:**
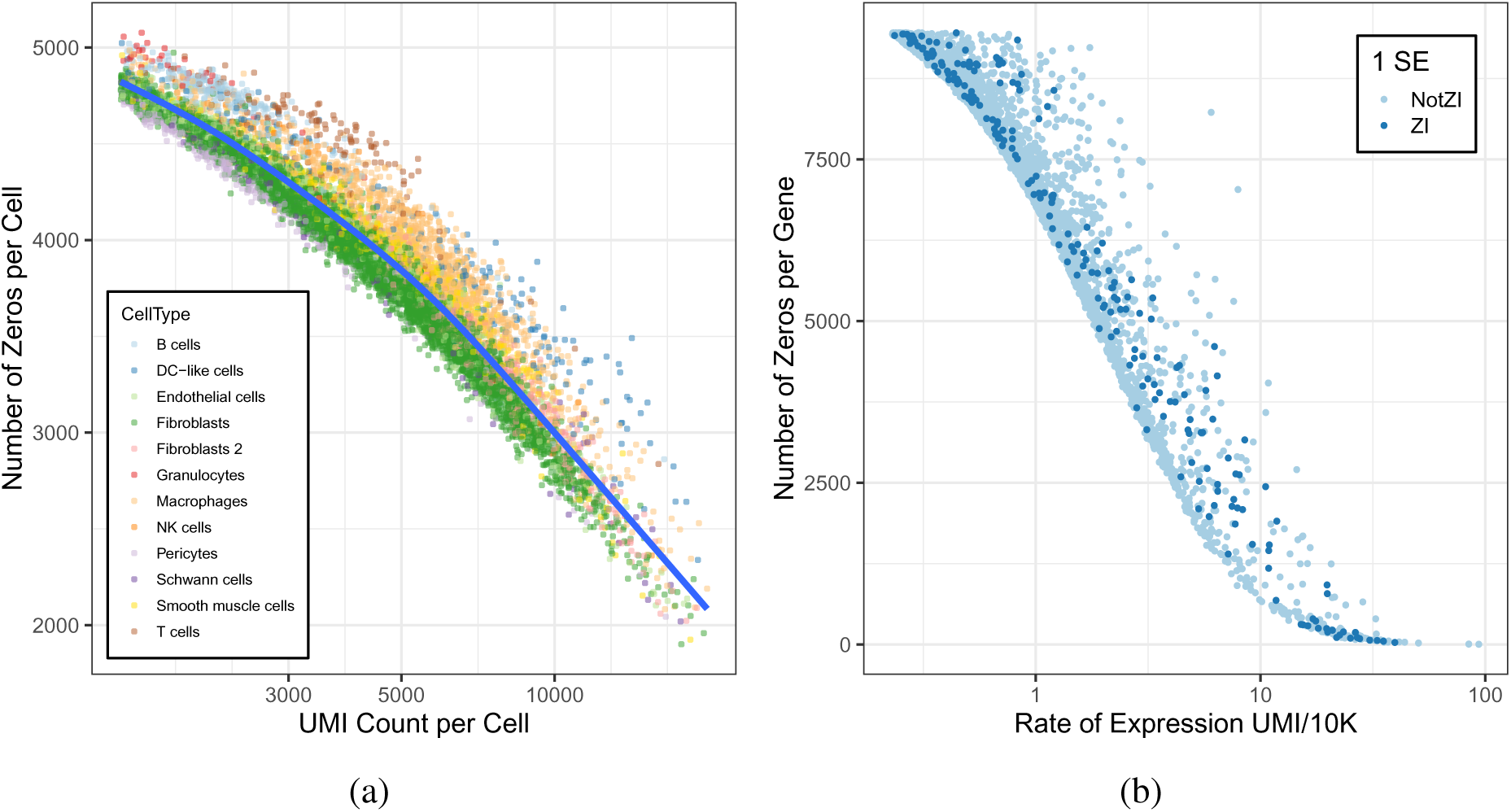
Factors that determine the number of zeros in scRNA-Seq data. (a) Total UMI counts per cell, which range from 746 to 17,302 with average 3,819 UMIs per cell, are plotted against the number of zeros per cell. Color coding indicates the individual cell types as determined by data-driven clustering. The blue line is the loess fit to the combined data (*R*^2^ = 0.947). (b) The per-gene rates of expression (*µ_g_*), which range from 0.23 to 97.4 with average 1.51 UMI/10K, are plotted against the number of zeros per gene. Genes that were identified as zero-inflated by `scRATE` (1 SE) are indicated in dark blue.

### Model selection can identify genes exhibiting zero inflation

In order to identify genes with zero inflation, we implemented a Bayesian model selection criterion— the expected log predictive density (ELPD) [Vehtari et al., 2017] — in our software package, `scRATE` (https://github.com/churchill-lab/scRATE). The ELPD score estimates out-of-sample predictive accuracy of four statistical models (P, ZIP, NB, or ZINB). It penalizes both underfitted and overfitted models. It examines all of the data, including non-zero counts, to provide a more complete evaluation of the count distributions than approaches that focus only on the zeros [Townes et al., 2019,Svensson, 2020]. `scRATE` uses leave-one-out cross-validation, which provides a standard error (SE) to quantify uncertainty in the estimated ELPD scores. The four models being compared have varying levels of complexity (P*≺*NB, P*≺*ZIP, NB*≺*ZINB, and ZIP*≺*ZINB) and in order to ensure that a more complex model is selected only when the ELPD is substantially better, we require that the difference in ELPD between two models is greater than zero by a multiple of the SE (e.g., 0 SE, 1 SE, 2 SE, and 3 SE). In addition to the model selection criterion, `scRATE` reports Bayesian parameter estimation and it can be used as a replacement for or as a complementary analysis tool along with standard GLM software.

To evaluate the true positive and false positive rates for detecting zero-inflated (ZI) genes — genes for which either the ZIP or ZINB model is selected — we simulated data similar to the heart data but with fixed levels of zero inflation (̂*π*) ranging from 0 to 90% and depth of sequencing at 10,000 UMIs/cell (*Simulation I* in Methods and Supplemental FigureS2a). We applied `scRATE` to the simulated data using the 0, 1, and 2 SE thresholds (Table1). `scRATE` has a high false positive rate at 0 SE but at the 1 SE threshold, the false positive error rate falls below 0.05 and at the 2 SE threshold, false positives are controlled at a stringency suitable for multiple testing across genes. In addition, we simulated data with average sequencing depths up to 50,000 UMIs/cell — higher than most droplet scRNA-Seq data — and observed a substantial improvement in power (Supplemental FigureS2b). This suggests that deeper coverage may be beneficial for detecting ZI genes. We carried out additional simulations to examine performance of different thresholds as described in Methods (*Simulation II*) and Supplemental Materials (Supplemental FigureS3). Having established that `scRATE` can detect ZI genes in simulated data, we next applied our model selection criterion to the heart data.

### scRNA-Seq data are zero-inflated for some genes

Townes [Townes et al., 2019] and Svensson [Svensson, 2020] have shown that the P or NB models, without zero inflation, are sufficient to capture technical variability of scRNA-Seq data. It is still of interest to determine whether these models are flexible enough to also capture biological hetero-geneity. In order to evaluate whether and how biological factors are contributing to zero inflation, we initially analyzed the heart data without considering any associated biological knowledge. We applied `scRATE` to each of 5,515 genes to classify them according to their best fitting model (Table2a) and to identify ZI genes. We found that for 1,474 genes, the best model (0 SE) is one of the ZI options. Using more conservative thresholds we found 220 genes (1 SE), 76 genes (2 SE), or 35 genes (3 SE) were best fit by a ZI model.

In order to evaluate the extent of under-calling of ZI genes by `scRATE`, we first down-sampled the data by randomly selecting subsets of cells and then repeated the model selection analysis (*Simulation III* in Methods and Supplement FigureS6). At the 0 SE and 1 SE thresholds, the numbers of ZI genes detected approaches saturation at or below 10,000 cells. However, for the 2 SE or 3 SE thresholds, the number of ZI genes detected steadily increases with sample size. This indicates that the data are under-powered to detect ZI genes at the 2 SE threshold even with sample size >10,000 cells. The ZI genes detected at 2 SE represent a lower bound on the number of high-confidence ZI genes that might be detected in a larger number of cells.

It seems intuitive that ZI genes would have a higher proportion of zeros and lower average expression when compared to other genes [Kharchenko et al., 2014]. However, our findings support the opposite conclusion (Figure2a, Supplemental TableS1). In our analysis of the heart data, ZI genes often have a lower proportion of zeros and higher rates of expression compared to genes that are not ZI (Figure2b,2c, and 2d). In cells where ZI genes are expressed, they exhibit higher average levels of expression compared to genes without zero inflation. The proportion of zeros is on average higher for NB genes compared to ZINB genes. Thus, the number of zeros alone is not a good indicator of zero inflation; rather, one must consider the entire count distribution to establish that zero inflation is present.

**Figure 2:**
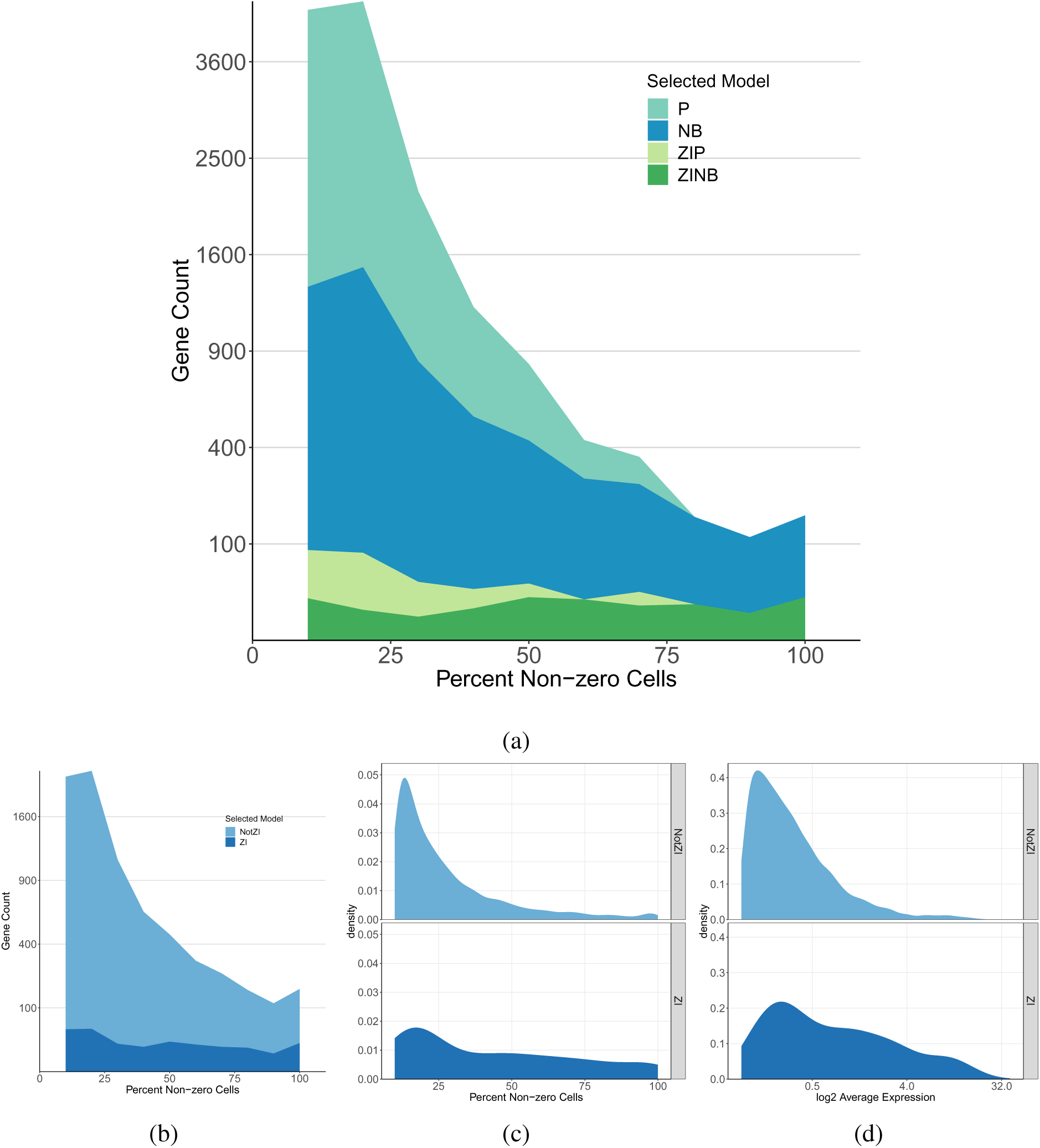
Classification of genes by `scRATE` using the threshold of 1 SE. (a) A density histogram shows the model selection for genes by `scRATE` as a function of percent non-zero cells. The ZI genes are uniformly distributed across the range, including genes with few zero counts. (b) Density histogram of `scRATE` classification collapsed to show only the ZI versus NotZI genes across percentages of non-zero cells. (c) Distribution of percent of cells with non-zero UMI counts for genes according to `scRATE` classification. (d) Distribution of average expression levels of genes according to `scRATE` classification. Also see Supplemental Figure S11 for the results with the other (0, 2, and 3 SE) thresholds.

### Most zero-inflated genes are due to variable expression rates across cell types

Skelly et al. [Skelly et al., 2018] classified each cell in the heart data into one of 12 cell types by data-driven clustering and integration of previous biological knowledge. The annotated cell types are heterogeneous and include cell types that are similar to one another (e.g., macrophages and dendritic cells) and cell types that are very different (e.g., smooth muscle cells and B cells). If zero inflation is primarily due to technical dropout, we would expect to see zeros evenly distributed across cell types. When we examined the distribution of zeros across cells in the ZI genes, we found that they tended to cluster within certain cell types (Figure3). The rate of expression of a gene is a major factor driving the frequency of zeros and, for many genes, the rate of expression varies widely across cell types. This suggests that we should evaluate zero inflation after taking cell type-specific rates of expression into account.

**Figure 3:**
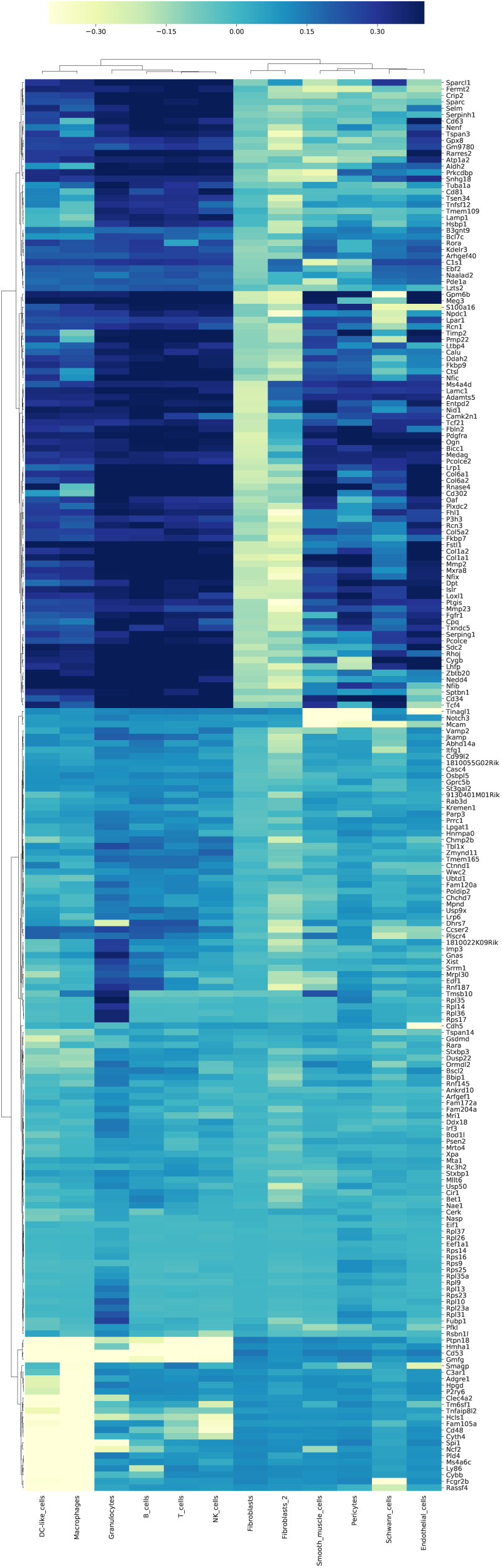
Zeros cluster within specific cell types. A bi-clustered heatmap of ZI genes (1 SE) by cell types shows the deviation from expected number of cells with zero UMI counts. Dark shading indicates an excess of zeros and light shading indicates that the cell type has fewer cells with zero UMI count than expected.

To account for biological variation in expression rates, we introduced cell type as an explanatory variable in the GLM and recomputed the `scRATE` classification (Table2b). After accounting for cell type, the number of zero-inflated genes drops markedly. Of the 76 genes that were originally classified as zero-inflated using the 2 SE thresholds, 72 are no longer classified as ZI, 3 genes remain ZI (*Xist, Rc3h2, Cir1*), 3 genes become ZI after accounting cell type (*Prnp, Folr2, Tax1bp2*), and for one gene (*Mmp2*) the `scRATE` algorithm failed to converge when cell type was included in the model. Genes that are no longer ZI after accounting for cell type display variation in rates of expression across cell types, such as *Col1a2* which is expressed primarily in fibroblasts, or *Ptpn18* which is expressed primarily in immune cells (Supplemental FigureS4).

In order to assess if this change in number of ZI genes was due to fitting the more complex model, we shuffled the cell type labels and repeated the `scRATE` classification. Results with the labels shuffled are similar to the `scRATE` classification without cell type (Table2c), demonstrating that the reduction in detected ZI genes is not due to a loss of power when including cell type as an explanatory variable.

While the majority of genes that were originally classified as ZI are no longer ZI after accounting for cell type, there are a handful of genes that remain or become ZI. Among them, *Xist* is an X chromosome silencing gene that is expected to be expressed only in female cells (Supplemental FigureS5). The heart data represent a mixture of female and male cells. We were able to unambiguously classify 63% of cells in silico as female or male in origin based on the presence of UMIs associated with female-specific *Xist* or with the Y-chromosome gene *Ddx3y*. `scRATE` classifies *Xist* as a zero-inflated gene at all thresholds up to 2 SE but *Ddx3y* is classified as NB and is only classified as a ZI gene at 0 SE after adjusting for cell type. After accounting for sex as an explanatory variable, in the subset of cells where we could establish sex, these genes are no longer ZI.

We fit a ZINB model to all genes and compared the estimated proportion of zero inflation (̂*π*_0_), with and without cell type in the GLM (Figure4a). For most genes, decreases. This is most after. For genes that remain ZI after accounting for cell type, there is little change in ̂*π*. For a handful of genes, including those that become ZI only after accounting for cell type, ̂*π*_0_ increases. These changes in ̂*π*_0_ are consistent with expectations from the model selection analysis. Genes with higher values of ̂*π*_0_ are more likely to be classified as ZI.

**Figure 4:**
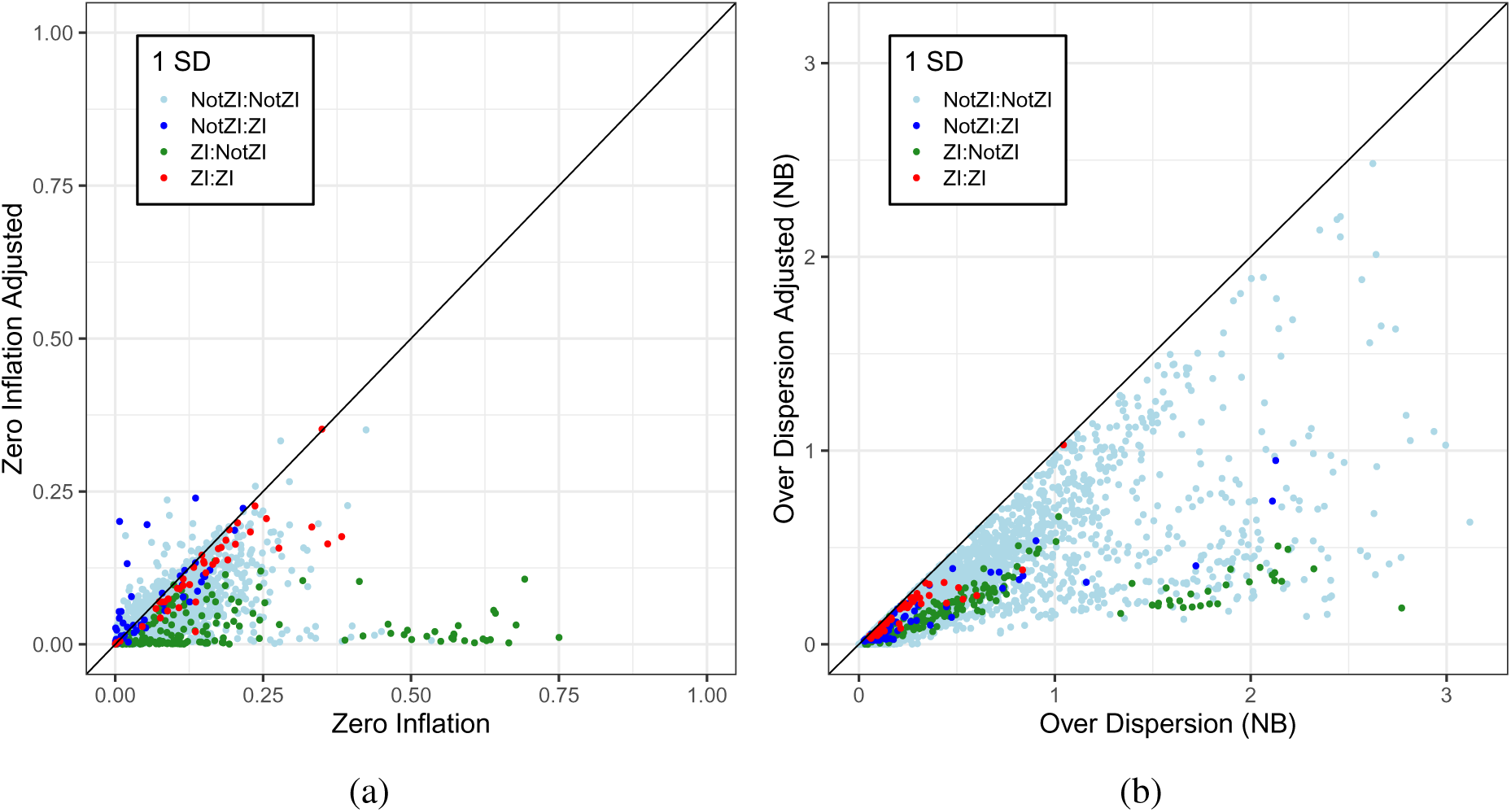
Effect of accounting for cell type on estimated zero inflation and overdispersion. (a) The scatterplot shows estimated zero inflation ̂*π* before and after including cell type in the GLM with ZINB error model. Color coding indicates the ZI classification of genes (1 SE) before and after accounting for cell type. The red point (ZI:ZI) at 0.3 on the diagonal is *Xist*. The light blue point (NotZI:NotZI) to the right is *Ddx3y*. (b) The scatterplot shows the estimated overdispersion *r*^ before and after including cell type in the GLM with NB error model.

Next we fit the NB model to all genes and compared the estimated overdispersion (∓) with and without cell type in the GLM (Figure4b). Accounting for cell type consistently reduces *r*^ and the effects on different classes of ZI genes are similar to those for the ̂*π*_0_ values. Thus, the overdispersion parameter of the NB model is able to identify much of the same heterogeneity that we are capturing with the ZINB model.

### Estimated zero inflation is not a reliable indicator of zero-inflated genes

The data features that distinguish the NB distribution from ZINB are subtle and, as a result, large sample sizes are needed to identify ZI genes (Supplemental FigureS2 and S6). It seems that we could avoid the problem of mis-classification by just fitting a ZINB model to each gene and reporting ̂*π*_0_ as a quantitative estimate of zero inflation. For example, for *Ddx3y*, after accounting for cell type, the estimated proportion of zero inflation is ̂*π*_0_ = 0.3326. This is comparable *Xist* for which ̂*π*_0_ = 0.3518. These sex-specific genes are genuinely zero-inflated (without accounting for sex) and although they are classified differently, the ̂*π*_0_ values are similar. The mis-classification inflation.

In order to evaluate the utility of ̂*π*_0_ as an indicator of zero inflation, we simulated NB and ZINB fit NB and ZINB models to each of the simulated datasets. We compared estimated values to the simulated truth (Supplemental FiguresS7,S8, and S9). Estimates of zero inflation from the ZINB model show a similar distribution for both the NB and ZINB simulated data (Figure5a,5b, and Supplemental FigureS9). We see that ̂*π*_0_ and can range as high as 50% for the NB simulated data, where the true value is zero. For the ZINB simulated data, ̂*π*_0_ is only weakly correlated with the simulated true ̂*π*_0_ (Figure5c and 5d). Our evaluation of suggests that it is not a reliable indicator of zero inflation.

**Figure 5:**
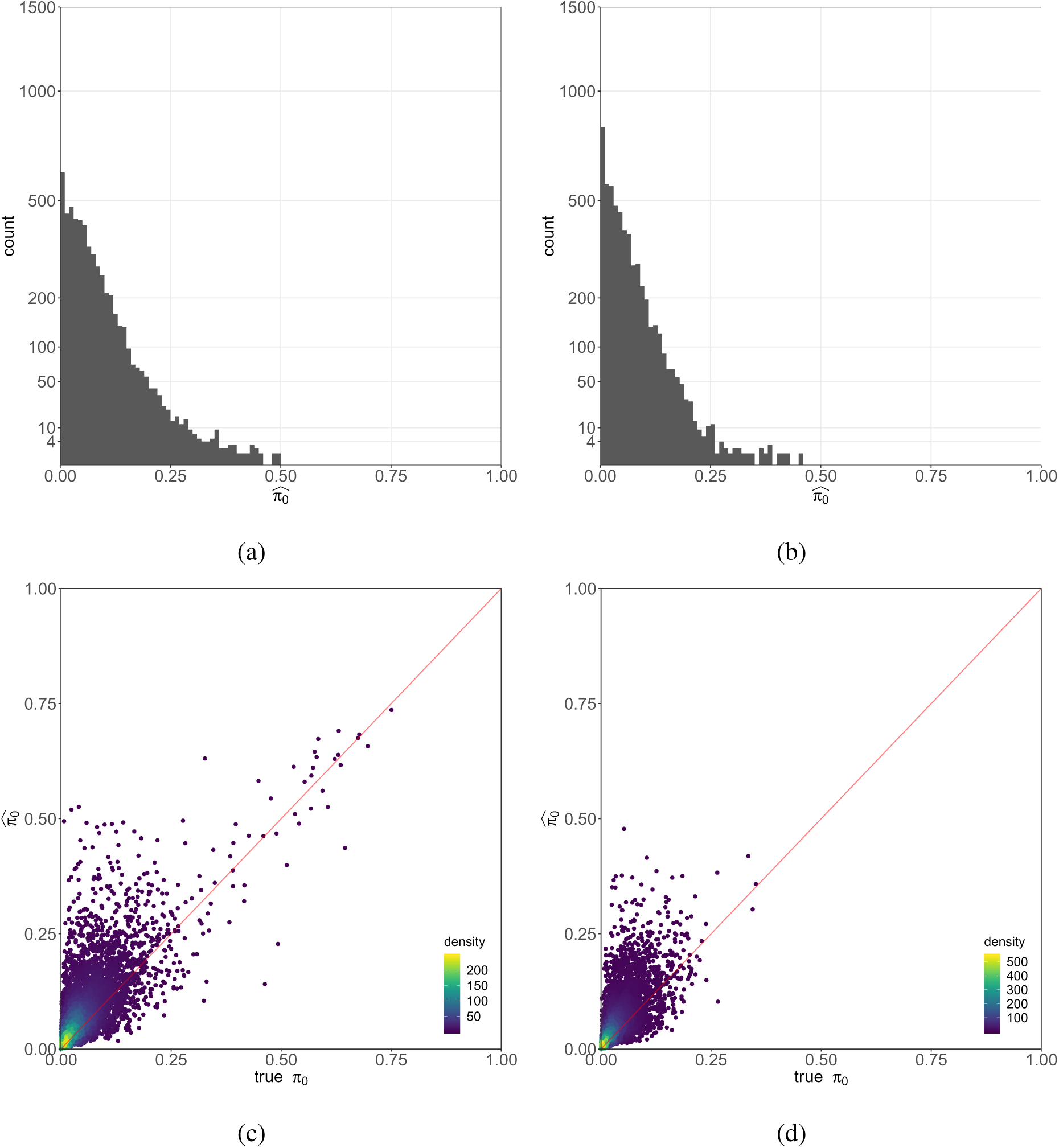
Estimating zero inflation with a ZINB model. Zero inflation probability ̂*π* estimated by ZINB on simulated NB data before cell type adjustment (a), and after cell type adjustment (b). Since simulated NB data does not contain zero inflation, it is implicit that the ZINB model should produce estimates of ̂*π* that are zero or very small. However, we find substantial overestimation of this quantity for many simulated genes. Scatter plots of true versus estimated zero inflation ̂*π*_0_ by ZINB on simulated ZINB data before cell type adjustment (c), and after cell type adjustment (d). Once cell type heterogeneity is regressed out, zero inflation is reduced.

## Discussion

Single cell RNA sequencing data display a high frequency of zero counts. The implications of this depend on understanding the processes that give rise to zeros. Looking across the entirety of cells in an experiment, we find that a substantial number of genes meet statistical criteria for zero inflation. However, this does not necessarily imply the existence of an independent zero-generating process such as technical dropout. Instead, we find that zero inflation is largely explained by biological factors, such as cell type and sex. Recent studies of scRNA-Seq in homogeneous cell populations confirm that there is no need to invoke technical dropouts as an explanation for zeros [Townes et al., 2019,Svensson, 2020].

A UMI count of zero does not necessarily imply that the gene is not expressed. This has led some researchers to propose imputation methods that convert zeros to non-zero values before analysis. We contend that zeros are informative data that should be incorporated directly into inferences about rates of expression and other parameters without modification. Based on the findings in this study we and others [Andrews and Hemberg, 2019,Townes et al., 2019,Svensson, 2020] recommend against the practice of replacing zeros in data with imputed non-zero values as this could potentially bias estimates of gene expression, reduce signatures of stochasticity, and mask biologically relevant heterogeneity.

Model selection criteria are useful for demonstrating the presence of zero inflation but we recommend against using a classifier to select gene-specific models for downstream analysis. This practice is known to result in inflated type-I error rates, especially when the power to discriminate among models is low [Gelman and Loken, 2014,Campbell, 2019]. Model averaging is one possible solution, but it can be computationally demanding and does not guarantee clear interpretation of parameters from models that we have averaged [Hooten and Hefley, 2019]. An alternative is to use a single, robust model that leads to reasonable inferences even when mis-specified. In our evaluation of the NB and ZINB models, the NB model produces accurate estimates of the mean and variance of gene expression across cells, even when applied to ZINB simulated data (Supplementary FiguresS7 and S8). Moreover, the NB dispersion parameter ( *r*) is a good indicator of heterogeneity (
Figure4b and Supplemental FigureS10). There is no perfect model and while the ZINB model is attractive for its generality, our simulation studies (*Simulation IV*) indicate that it may not provide reliable inferences. We recommend the generalized linear model with negative binomial errors, an offset to account for cell-to-cell variation in depth of sequencing, and including known biological factors as explanatory variables. While there are certainly opportunities to improve on this analysis model, it serves as the logical baseline for evaluation of alternative approaches and refinements.

We identified cell type as a major contributor to heterogeneity in gene expression that can explain apparent zero inflation. Data-driven clustering is not always successful in delineating cell subtypes, and depends in part on the comparisons of interest to the analyst as well as the resolution with which data are viewed. Residual biological heterogeneity within a particular cell type classification may reflect distinct subgroups of cells, transient cell states, or variation along a continuum. Clustering analysis divides cells into discrete groups but cell types are often hierarchical and distinct clusters may share different degrees of similarity [Zappia and Oshlack, 2018]. Moreover, in some cases cell “types” may exist along a continuum [Stanley et al., 2019], making cluster boundaries somewhat arbitrary and dependent on features of the clustering algorithm and data. Persistence of zero inflation or high levels of over dispersion after accounting for cell type are indicators of unknown sources of biological variation that may prove to be useful in refining cell type hierarchies or positioning cells along the trajectories of a continuum.

In summary, we find substantial evidence for zero inflation in scRNA-Seq data, much of which can be explained by known biological factors including cell type and sex. There remain a number of ZI genes for which we have not identified a biological explanation. Genes with zero inflation can potentially help to reveal hidden biological factors such as stage in the cell cycle, activation status of immune cells, or incomplete classification of cell types that vary across the heterogeneous mixture of cells. The model selection procedure implemented in `scRATE` software provides an exploratory data analysis tool for identifying these interesting genes.

## Methods

### Data

The heart data [Skelly et al., 2018] consist of metabolically active, nucleated, non-myocyte cells from heart ventricles of female and male C57BL/6J mice. The dataset was sequenced on 10X Chromium scRNA-Seq platform. We used the preprocessed UMI counts (downloaded from https://www.ebi.ac.uk/arrayexpress/experiments/E-MTAB-6173/), originally obtained using cellranger version 1.3 (10X Genomics). Downstream analysis using Seurat version 2.0.0 [Butler et al., 2018,Macosko et al., 2015] identified 12 cell types over 10,519 cells. In order to ensure that we include only expressed genes in our analysis, we restricted attention to 5,515 genes that had at least 1 UMI in at least 10% of cells.

### Generalized linear models for count data

`scRATE` implements Bayesian estimation and model selection for generalized linear models (GLM) with or without zero inflation. The distribution of counts is modeled using the log link function as a linear combination of an offset and covariates. The effect of including the offset is to account for differences in total exposure (total UMI counts per cell). With the offset, the regression parameter estimates are scaled as rates of expression in units of UMI counts per 10,000. Including a categorical covariate, e.g., cell type, allows the rates to vary across groups of cells. The zero-inflated models include a second component with zero inflation parameter ̂*π*_0_ that represents the probability that an observed datum is an obligate zero. This component uses a logistic link function and does not require an offset. The expected number of zeros will be greater in cell types with lower rates of expression, but the proportions of extra zeros is constant across cells. Standard errors of estimated parameters are obtained by Monte Carlo sampling (`scRATE`) or by application of the robust sandwich estimator (`CountReg` [Zeileis et al., 2008]).

### Model selection

For counts associated with a given gene *y_c_*, where *c* is an index over cells, we fit Poisson, Negative Binomial, Zero-Inflated Poisson, and Zero-Inflated Negative Binomial models and evaluate their predictive accuracy based on Expected Log Predictive Density,

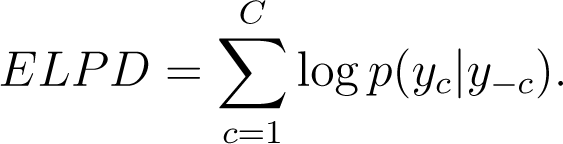

As a general rule, we select a model that has the largest mean ELPD as the best fit. But the data features that distinguish the best model from other models are subtle for many genes. The full Bayesian implementation provides an estimate of the mean and standard error (SE) of *ELPD difference* between models. In case two models provide similarly good fit, we select a simpler model unless the credible interval of ELPD margin offered by the other (more complex) model is always positive.

### Software

We have implemented our model selection approach in open-source R package, `scRATE`, available at https://github.com/churchill-lab/scRATE with GPLv3 license. The package is built around `rstanarm` [Goodrich et al., 2020], `brms` [Bürkner, 2017,Bürkner, 2018], and `loo` [Vehtari et al., 2019] and has many additional features that facilitate the parallel analysis on PBS/torque or SLURM clusters.

### Simulations

#### Simulation I: ZINB genes with known levels of zero inflation

We evaluated the power for detecting ZI genes by simulating ZINB data with known zero inflation probability (̂*π*_0_) of 10, 20, 30, *· · ·*, up to 90% based on the mean and shape parameters estimated from the mouse heart data. We generated data with two different sequencing depths of 10,000 and 50,000 UMIs/cell. As sequencing depth increases, we find model under-calling substantially reduces and the proportion of correct zero inflation calls increases (Supplemental FigureS2). This implies zero inflation is harder to detect when the sequencing depth is lower. We find many studies are performed below 10,000 UMIs/cell, for example, the median depth of coverage for the heart data was *∼*2,500 UMIs/cell.

#### Simulation II: Simulated genes with a mix of known distributions

Using model classification and parameters estimated from the heart data, we simulated P, NB, ZIP, and ZINB data for each of 5,515 genes. We simulated data based on the 0 SE classification by selecting distributions for genes according to the model called at 0 SE. We repeated this process using model calls at 1 SE, 2 SE, and 3 SE. We applied `scRATE` model selection with the 0, 1, 2 and 2 SE thresholds to each simulated gene set. These model calls allow us to compute true and false positive rates for detecting ZI genes and to compute the AUC for each combination of simulation and evaluation thresholds (Supplemental FigureS3). We find that the 1 SE threshold provides the best balance between false positive (over-calling of model) and false negative (under-calling of model) classification.

#### Simulation III: Random subsets of cells

The number of cells may have a substantial impact on the power of detecting ZI genes. In order to assess the effect of cell number on detecting ZI genes, we generated random subsets of cells by down-sampling data from the 10,519 cells in the mouse heart data. The number of cells ranges from 44 up to 9,000 (Supplemental FigureS6). These random subsets retain the heterogeneity of original data, and therefore, the number of ZI genes should not appreciably change with the number of cells, except due to loss of power for detecting ZI genes. We found that the number of ZI genes increases with the number of cells sampled (Supplemental FigureS6). This implies the number of ZI genes we detected in the original dataset should be regarded as a lower bound.

#### Simulation IV: NB versus ZINB

To evaluate the effect of naïvely applying a ZINB model to non-zero-inflated data, we simulated 5,515 genes from an NB distribution with parameters estimated from the heart data. We fit both NB and ZINB models to simulated NB data and evaluated the parameter estimates including ̂*π*_0_ for each gene. We find that fitting NB data with the ZINB model yielded high estimates of zero inflation for many genes. Next we simulated 5,515 genes from a ZINB distribution with parameters estimated from the heart data. We fit both NB and ZINB models to the simulated ZINB data. ZINB is the correct model in this simulation, but we find that ̂*π*_0_ estimation is still unstable for many TableS2). We also find that NB leads to reasonable inferences even when mis-specified in this simulation.

## Supplemental Materials: Bayesian model selection reveals biological origins of zero inflation in single-cell transcriptomics

### Additional Dataset I: the mouse kidney data

This dataset [Park et al., 2018] is a single cell transcriptome atlas of mouse kidney. The library was prepared with the whole kidney tissues of seven healthy male C57BL/6 mice using 10X Chromium (v2 chemistry) protocol and sequenced on Illumina HiSeq 2000 platform. We downloaded a processed data file https://www.ncbi.nlm.nih.gov/geo/download/?acc=GSE107585&format=file&file=GSE107585_Mouse_kidney_single_cell_datamatrix.txt.gz available at NCBI’s GEO under accession number GSE107575. It includes 43,745 cells consisting of 16 distinct cell types: endothelial, vascular, descending loop of Henle, podocyte, proximal tubule, ascending loop of Henle, distal convoluted tubule, collecting duct (CD) principal cell, CD intercalated cell, CD transitional cell, fibroblast, macrophage, neutrophil, natural killer cell in addition to two novel cell types. In order to ensure that we include only expressed genes in our analysis, we restricted attention to 5,160 genes that had at least 1 UMI in at least 10% of cells, the same filtering criterion used for the heart data.

### Additional Dataset II: the human PBMC data

This dataset [10X Genomics, 2018] is peripheral blood mononuclear cells (PBMCs) from a healthy donor, sequenced on Illumina NovaSeq platform with ∼ 54,000 reads per cell. We downloaded a processed data file [Stuart et al., 2019] available at https://www.dropbox.com/s/zn6khirjafoyyxl/pbmc_10k_v3.rds? dl=0. It contains 9,432 cells consisting of 14 cell types: CD14+ Monocytes, CD4 Memory, CD4 Naive, pre-B cell, Double negative T cell, NK cell (bright and dim), B cell progenitor, CD8 effector, CD8 Naive, CD16+ Monocytes, Dendritic cell, pDC, Platelet. The raw data are also available at https://support.10xgenomics.com/single-cell-gene-expression/datasets/3.0.0/pbmc_10k_v3. In order to ensure that we include only expressed genes in our analysis, we restricted attention to 6,435 genes that had at least 1 UMI in at least 10% of cells, the same filtering criterion used for the heart data.

---

**Supplemental Figure S1:**
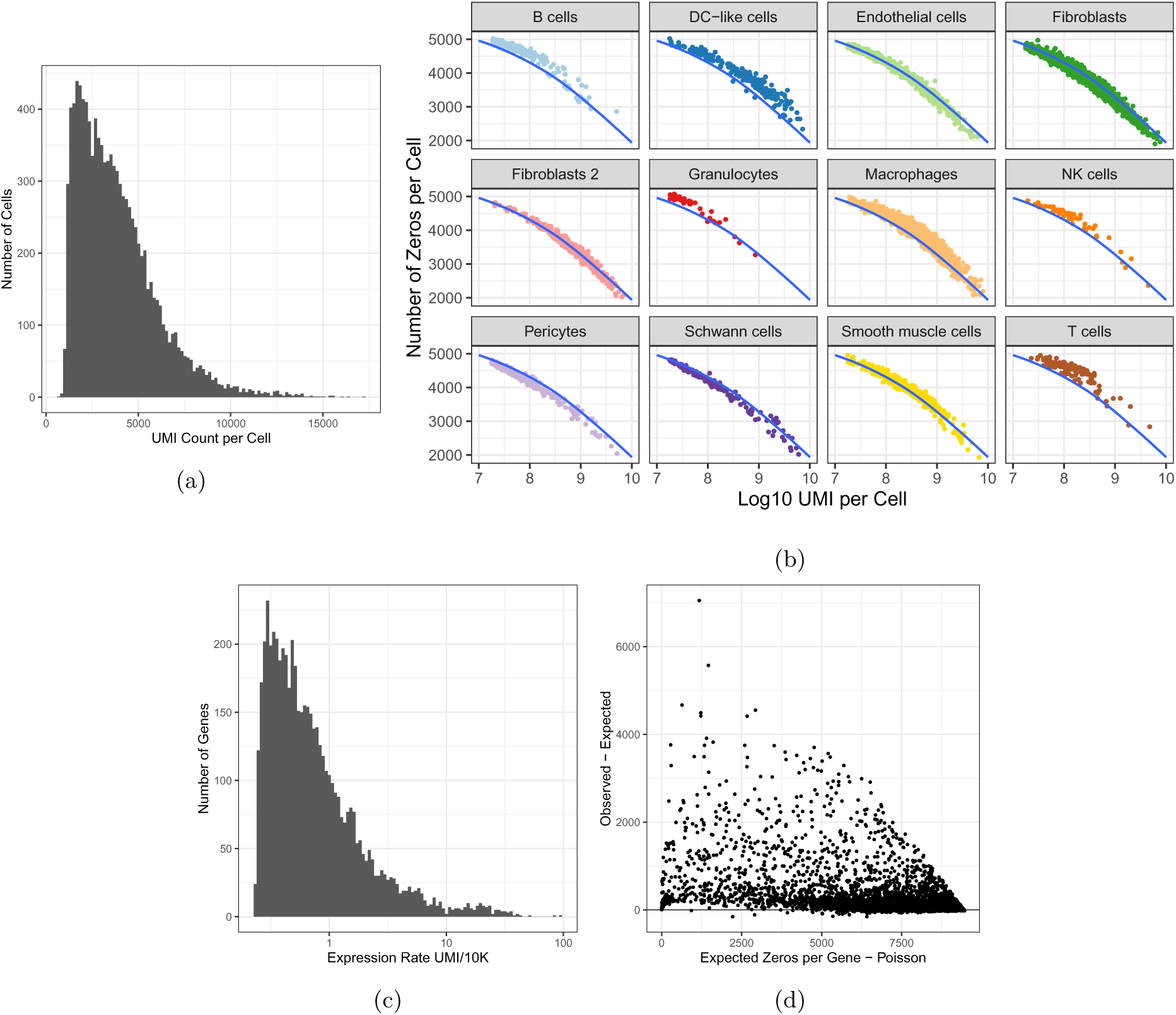
Factors that determine the number of zeros in scRNA-Seq data. (a) Total UMI counts per cell, which range from 746 to 17,302 with average 3,819 UMIs per cell, are shown as a histogram. (b) The number zeros per cell is plotted against the log10 UMI count. The plot is facetted to show the individual cell types as determined by data-driven clustering. The blue line shows the loess fit to the combined data. (c) The per-gene rates of expression (*µ_g_*), which range from 0.23 to 97.4 with average 1.51 UMI/10K, are shown as a histogram. (d) A scatter plot shows the expected number of zeros under Poisson sampling (x-axis) compared to the difference from expectation (observed expected; y-axis) and reveals genes with an excess of zeros.

**Supplemental Figure S2:**
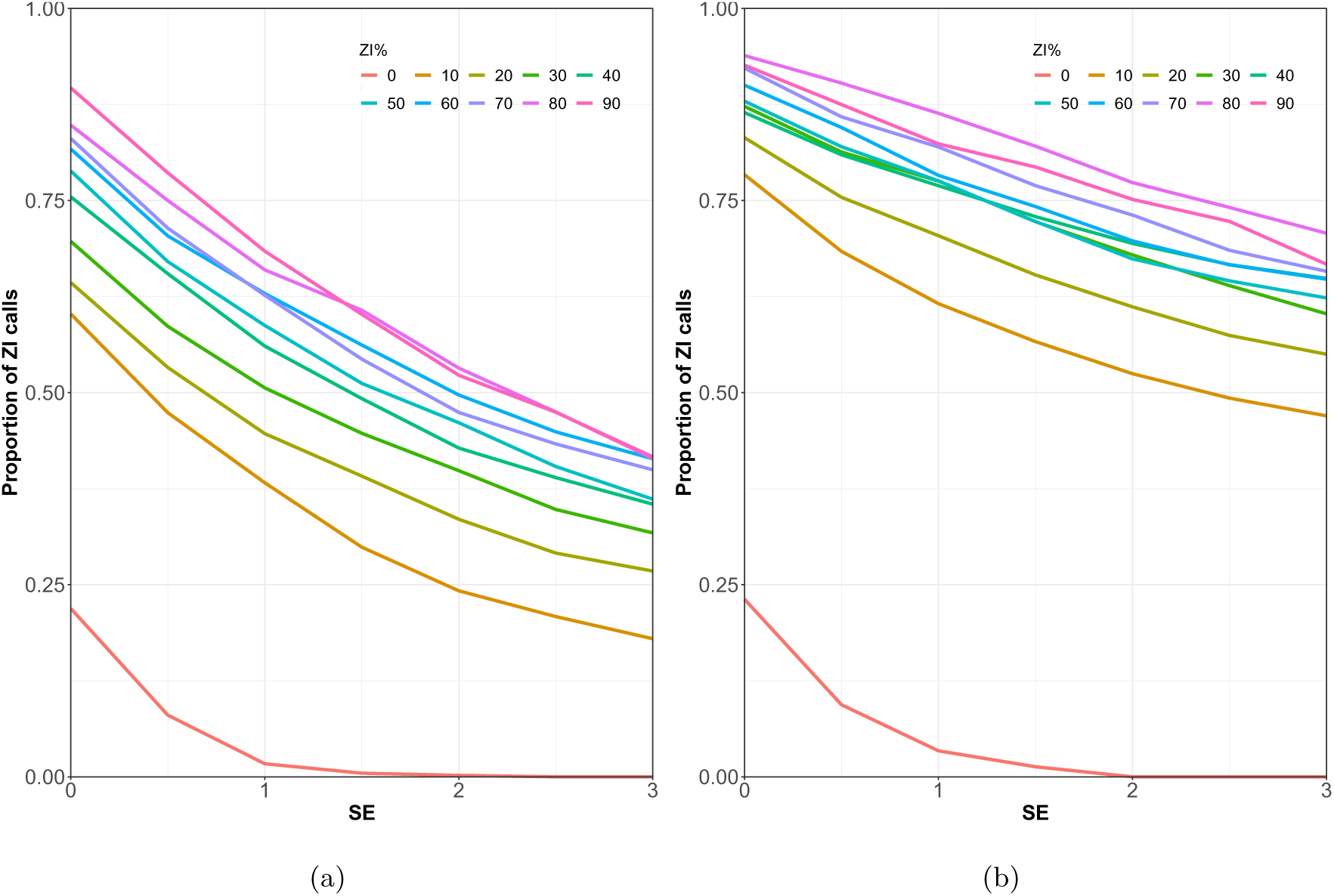
Power analysis of scRATE classification of ZI gene. We simulated data as described in Methods (*Simulation I*) across a range of zero inflation (0 to 90%, color coded). We applied scRATE with thresholds ranging from 0 SE to 3 SE (x-axis) and counted the proportion of genes classified and ZIP or ZINB (y-axis). The red line shows the numbers of ZI genes detected when the simulation model is NB, with no zero inflation. This is the false positive rate, or type-I error, of the classifier. The other lines indicate different proportion of zero inflation in the ZINB model. This is true positive rate, or power, of the classifier. Simulations are based on sequence depths of 10,000 UMIs (a) and 50,000 UMIs per cell (b).

**Supplemental Figure S3:**
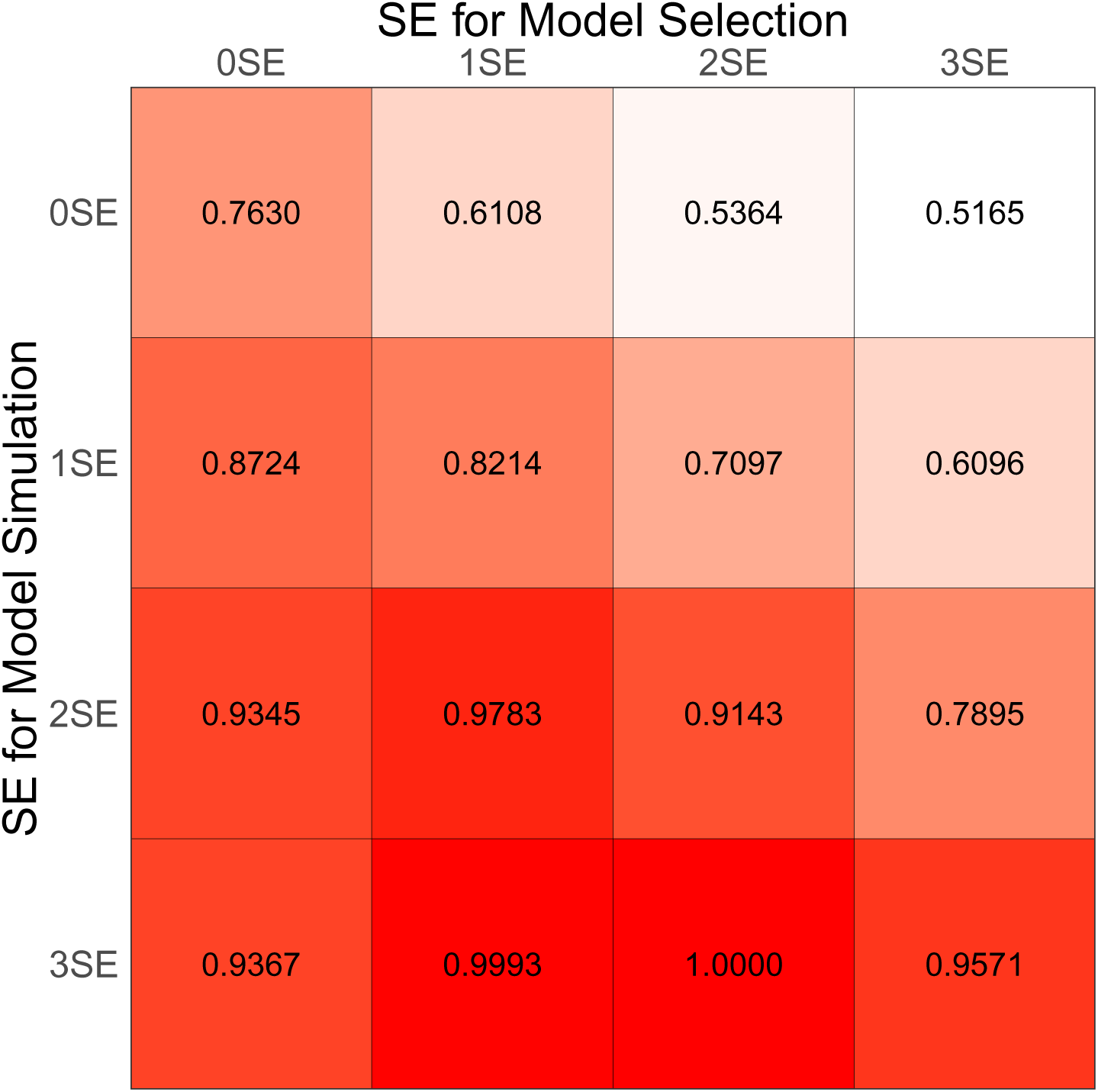
Area Under ROC Curve (AUC) of different model selection thresholds. As shown in Supplemental Figure S2, there is a trade off along the stringency of threshold: the more stringent it gets, the rate of model under-calling (false negative rate) increases while the rate of over-calling (false positive rate) reduces. In order to identify the optimal threshold, we evaluated AUC with simulated gene sets (*Simulation II*) for which distributions are selected from the heart data with the 0, 1, 2, and 3 SE thresholds (rows). For each simulated gene set, we performed scRATE classification with the 0, 1, 2, and 3 SE thresholds (columns). We find that the 1 SE threshold (the second column) is robust and performs relatively well across the simulated gene sets.

**Supplemental Figure S4:**
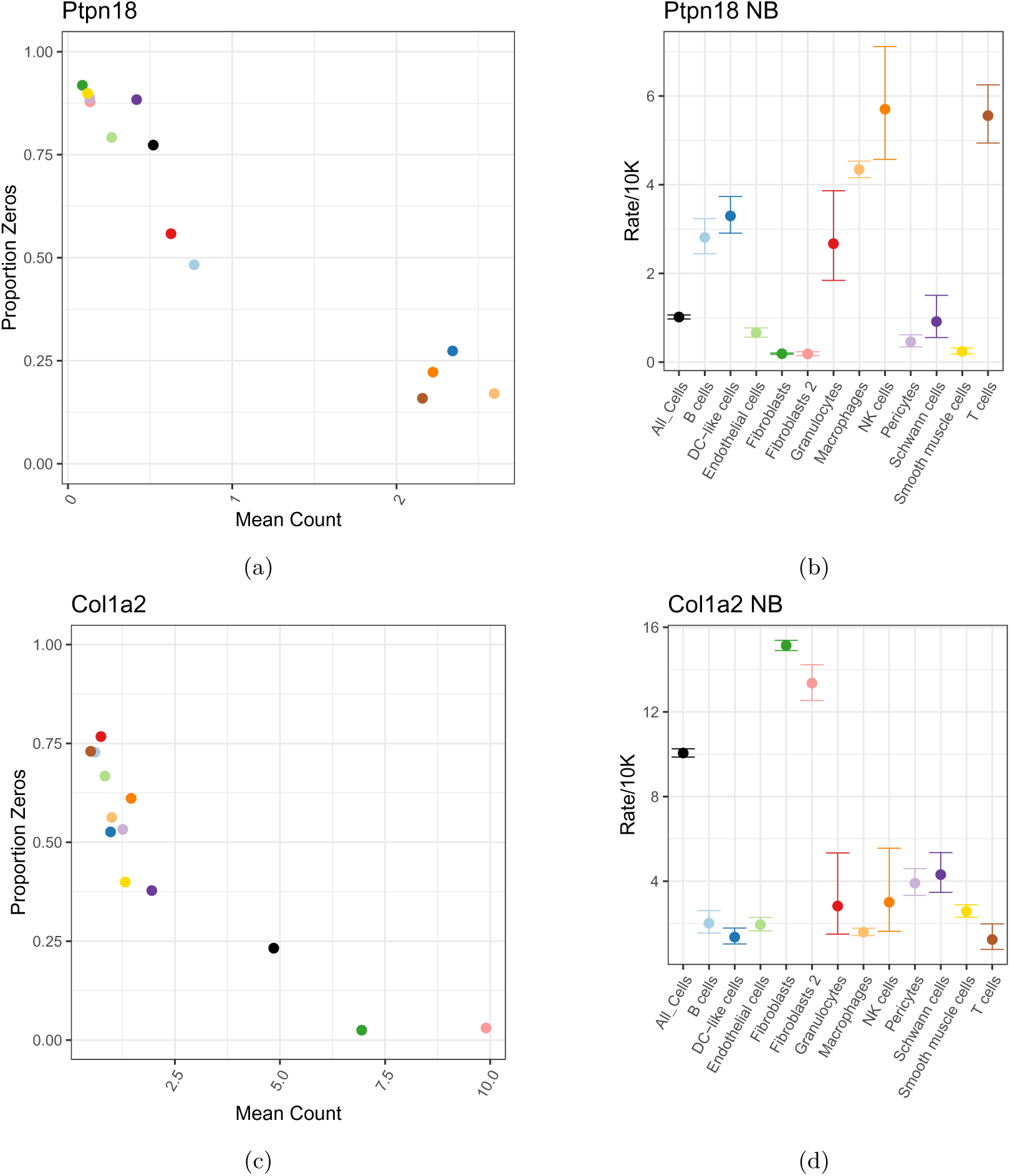
Examples of ZI genes that are no longer ZI after accounting for cell type. *Ptpn18* is primarily expressed in immune cells. The upper left panel shows the proportion of zeros, averaged across cells within each cell type, as a function of the mean UMI count per cell. The upper right panel shows the estimated rates of expressed for NB model overall (All Cells) and for each cell type as estimated by scRATE with cell type as covariate. Lower panels show the same for *Col1a2*, a gene primarily expressed in fibroblasts.

**Supplemental Figure S5:**
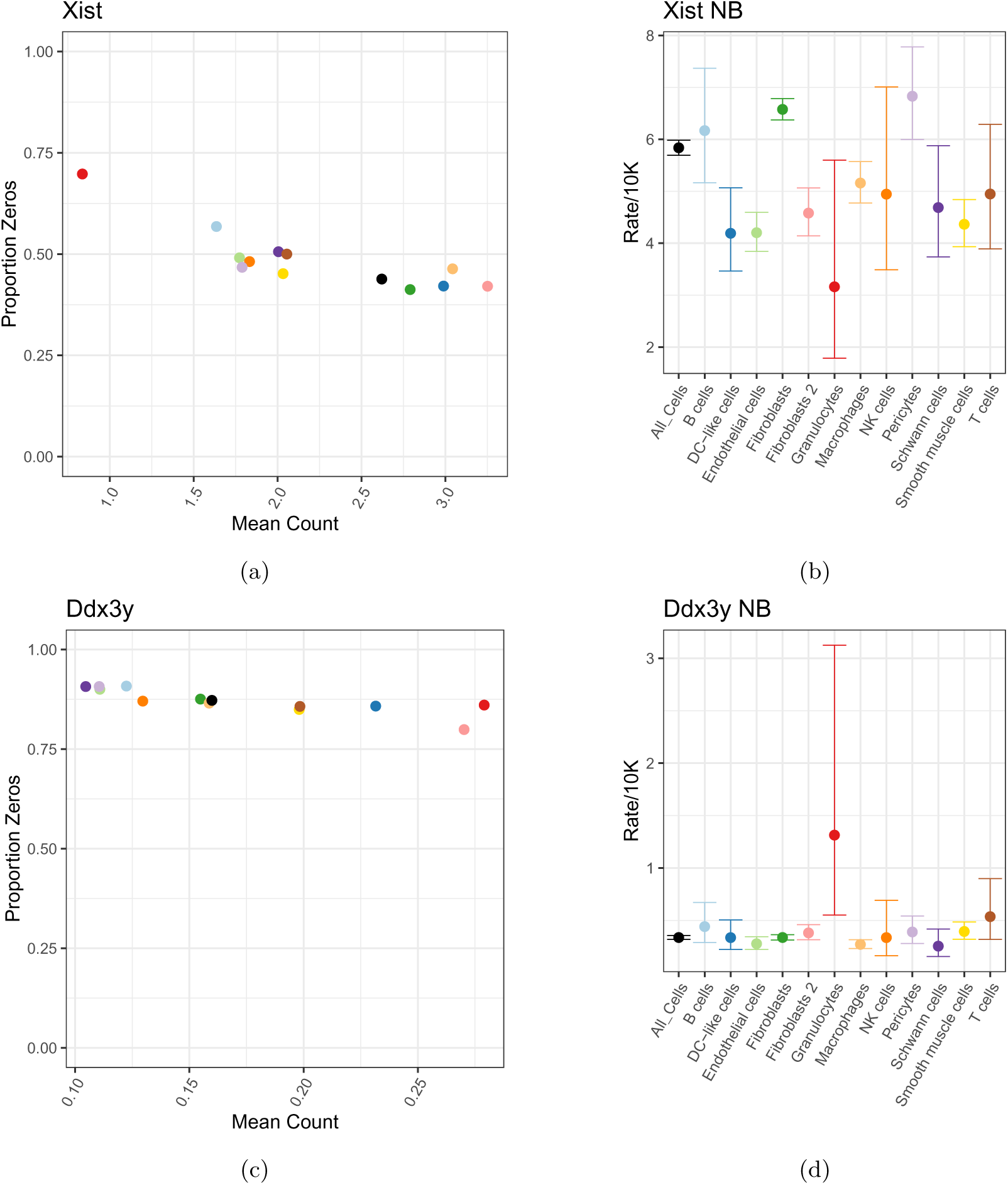
Examples of ZI genes that remain ZI after accounting for cell type. *Xist* is a female specific transcript encoded on the X chromosome. *Ddx3y* is a male specific gene encoded on the Y chromosome. Panels are as described for Supplemental Figure S4. There appears to be a high proportion of granulocytes among the male cells.

**Supplemental Figure S6:**
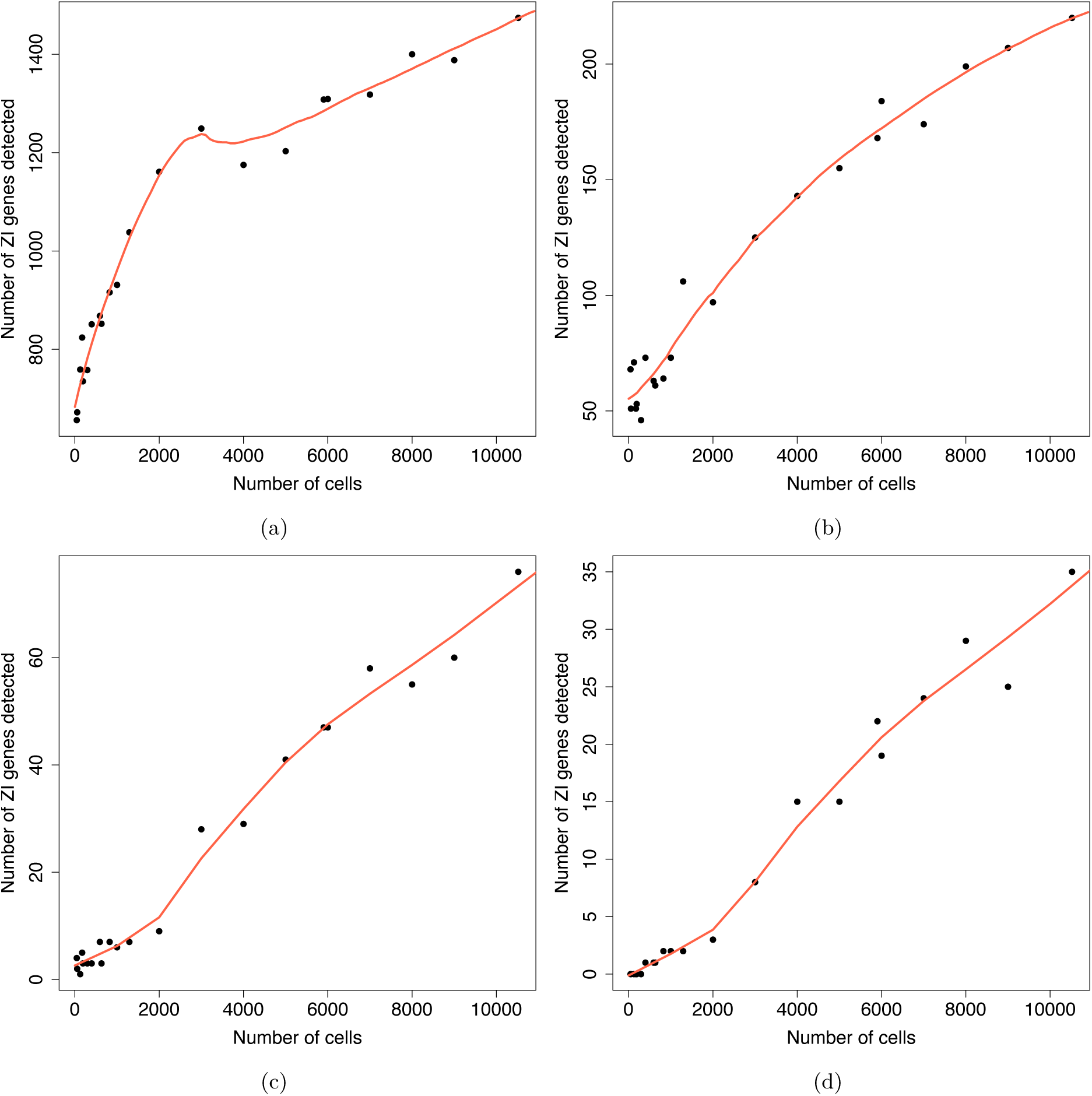
Detection of ZI genes in down-sampled heart data. We randomly sampled subset of cells (*Simulation III*) across a range of sample sizes (x-axis), applied scRATE to the reduced data, and counted the numbers of ZI genes (y-axis) at the threshold of 0 SE (a), 1 SE (b), 2 SE (c), and 3 SE (d). Red line is a lowess fit.

**Supplemental Figure S7:**
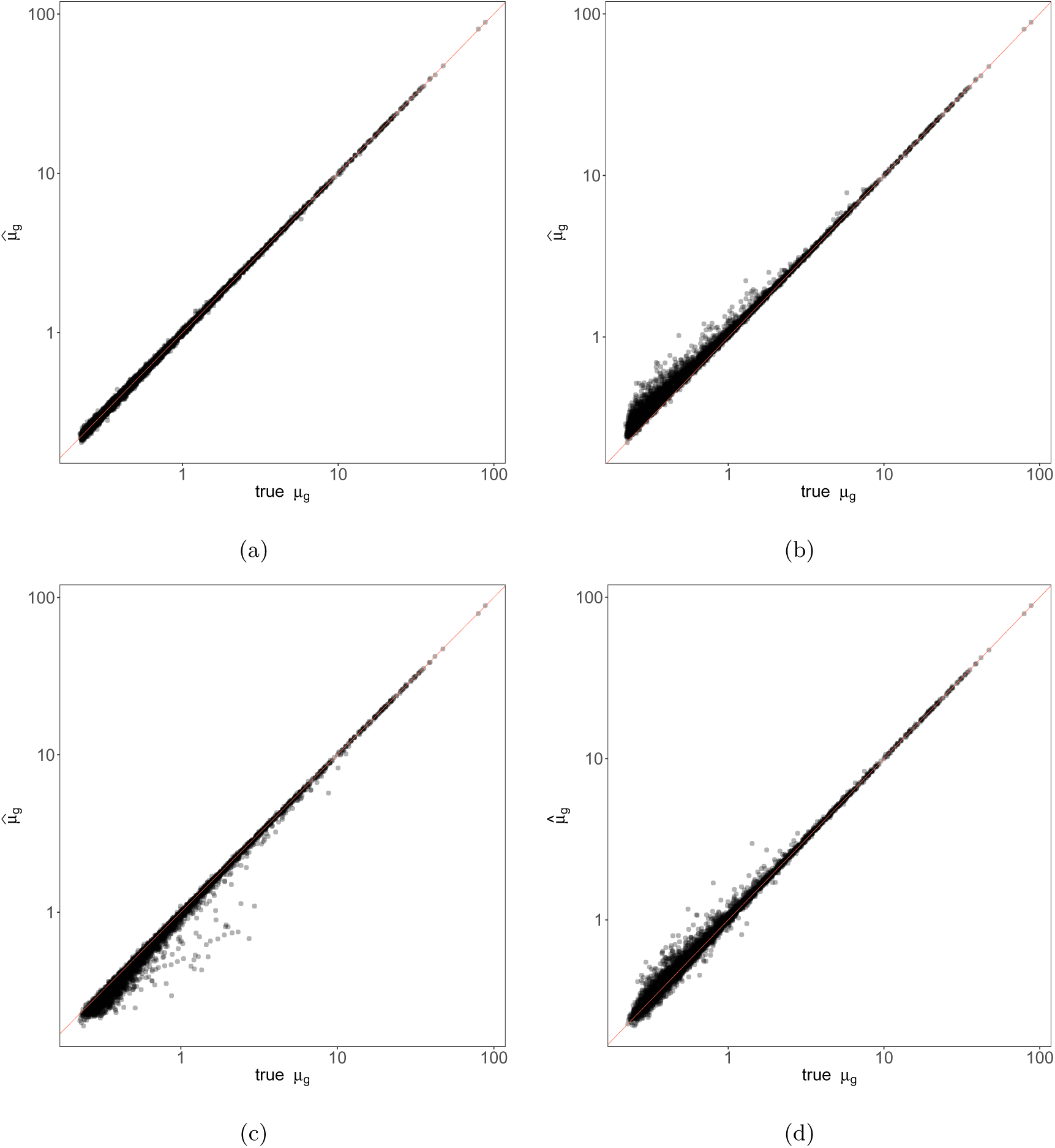
Estimation of the mean (rate of expression) parameter in simulated data. Data were simulated under either the NB model or ZINB model as described in Methods (*Simulation IV*). All panels show estimated rates (y-axis) are compared to simulation truth (x-axis). (a) Fitting NB model to NB simulated data. (b) Fitting ZINB model to NB simulated data. (c) Fitting NB model to ZINB simulated data. (d) Fitting ZINB model to ZINB simulated data. The biases observed in (b) and (c) are explained by the different interpretation of mean rate between the NB and ZINB models.

**Supplemental Figure S8:**
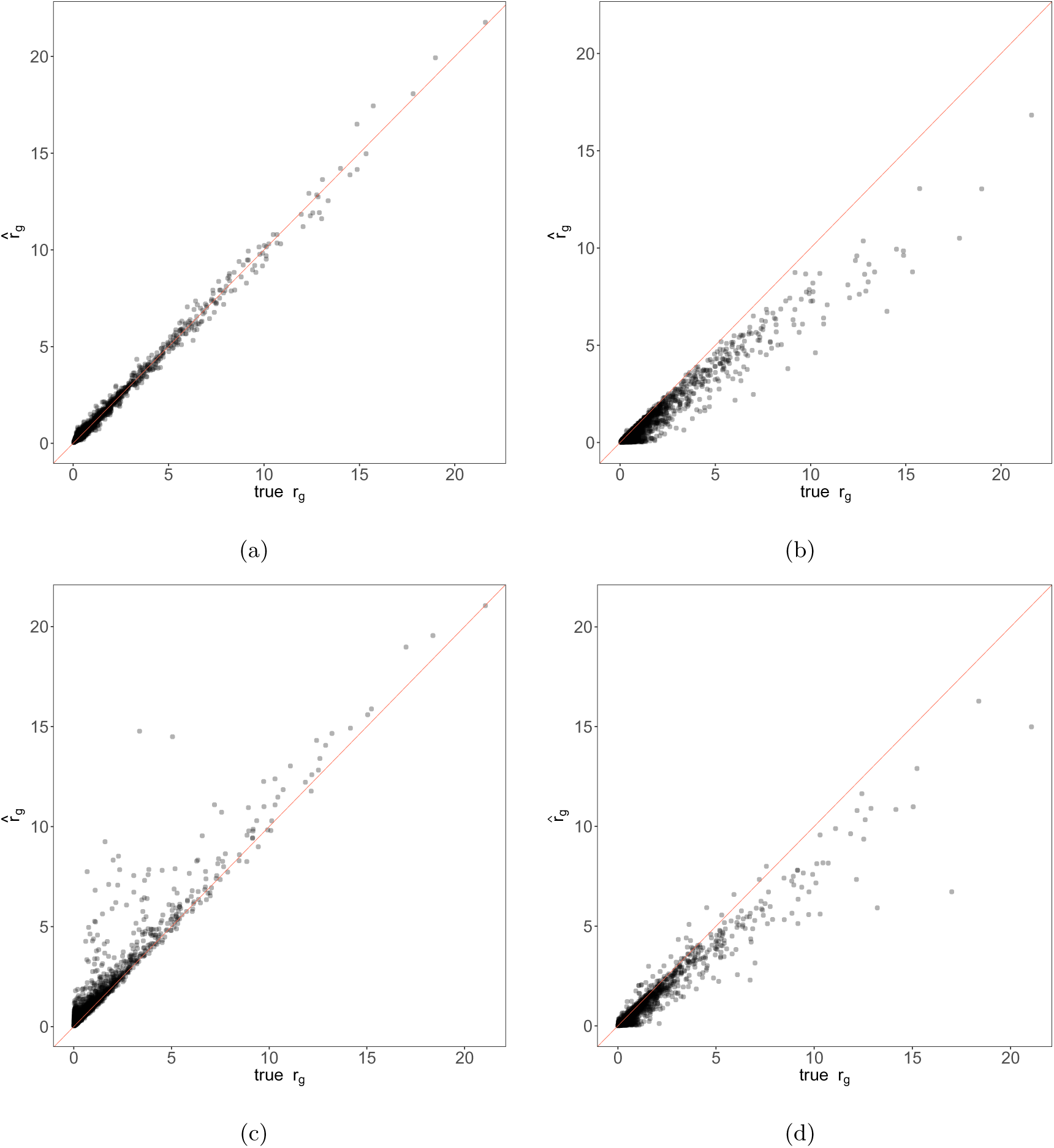
Estimation of the overdispersion in simulated data. Data were simulated under either the NB model or ZINB model as described in Methods (*Simulation IV*). All panels show estimated overdispersion (y-axis) are compared to simulation truth (x-axis). (a) Fitting NB model to NB simulated data. (b) Fitting ZINB model to NB simulated data. (c) Fitting NB model to ZINB simulated data. (d) Fitting ZINB model to ZINB simulated data. The biases observed in (b) and (c) illustrate how the overdispersion and zero inflation parameters trade-off, with one compensating for the other when fitted model is mis-specified relative to the data.

**Supplemental Figure S9:**
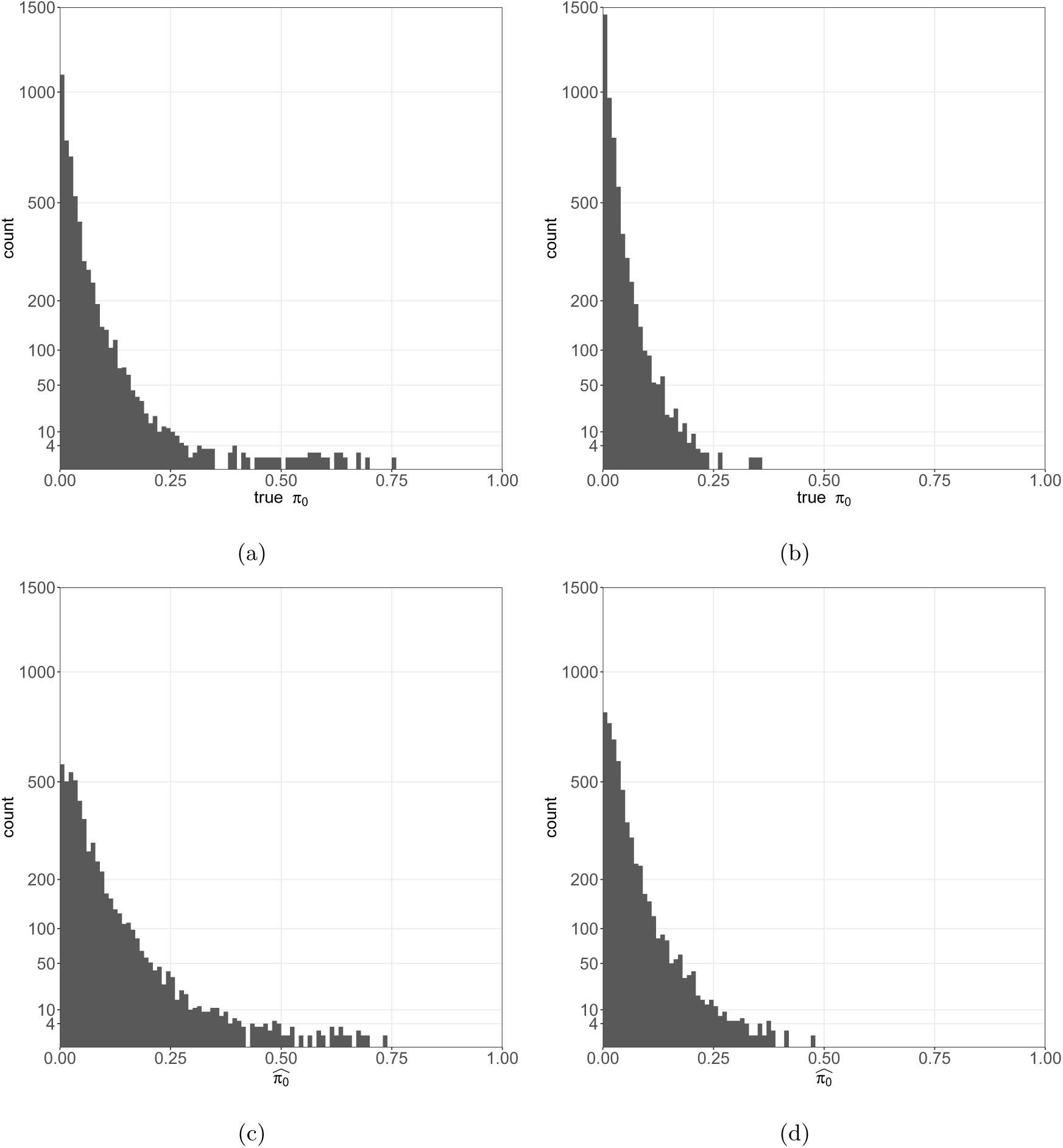
Estimated zero inflation on simulated ZINB data. True zero inflation ̂*π*_0_ estimated from the mouse heart data before cell type adjustment (a) and after cell type adjustment (b). Estimated zero inflation ̂*π* on simulated ZINB data before cell type adjustment (c) and after cell type adjustment (d). Cell type t reduces the estimated amount of zero inflation overall. The variability of estimated

**Supplemental Figure S10:**
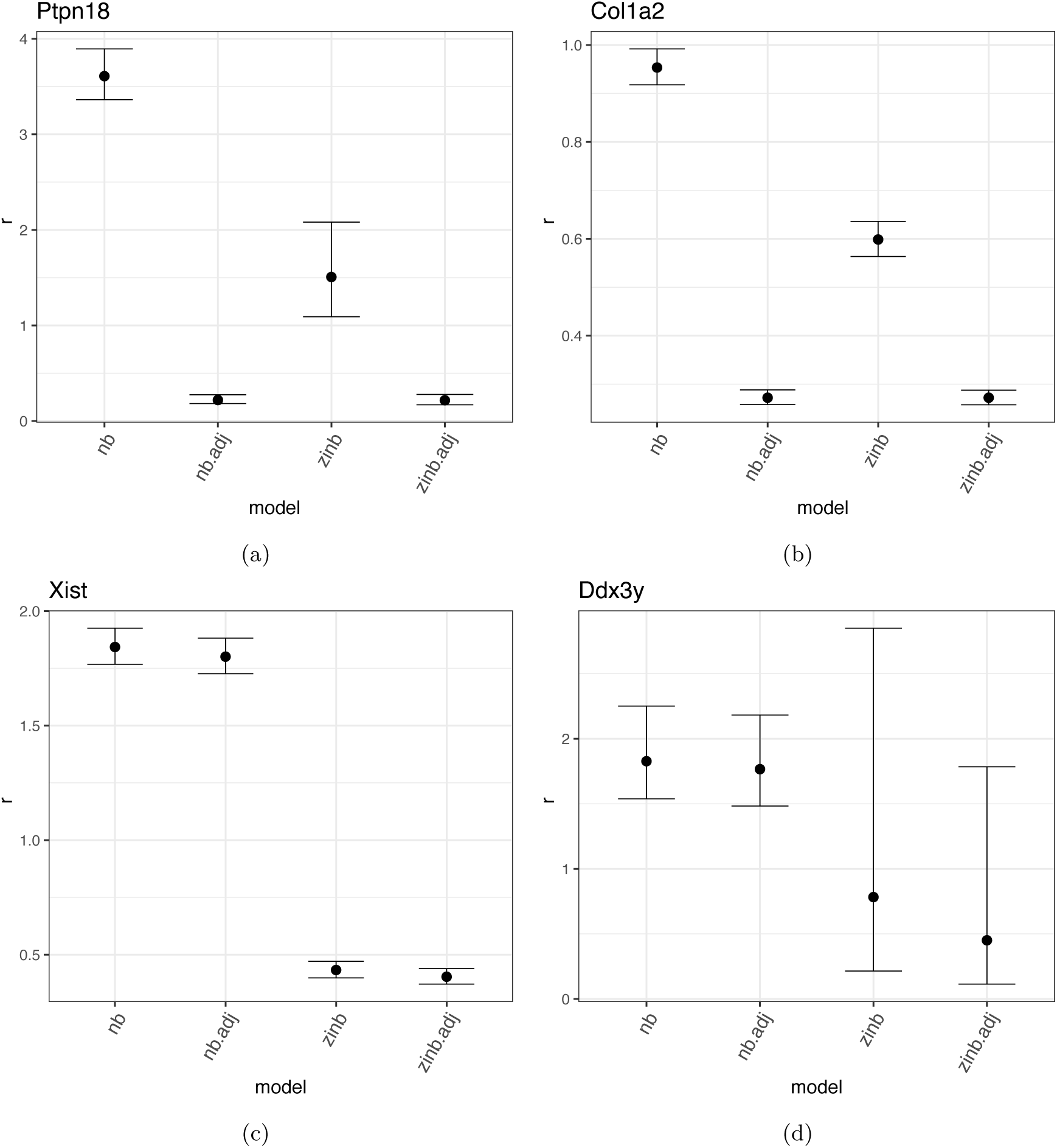
Effects of cell type and zero inflation on estimated overdispersion. The overdispersion parameter (y-axis) was estimated using the NB model, with and without cell type as a covariate and again using the ZINB model with and without cell type. For the cell type-specific genes *Ptpn18* and *Col1a2*, there is a small reduction in overdispersion from the NB to ZINB model without cell type. Including cell type as a covariate reduces overdispersion to almost zero. For the sex-specific genes *Xist* and *Ddx3y*, the inclusion of cell type has negligible effect on overdispersion whereas allowing for zero inflation reduces overdispersion substantially.

**Supplemental Figure S11:**
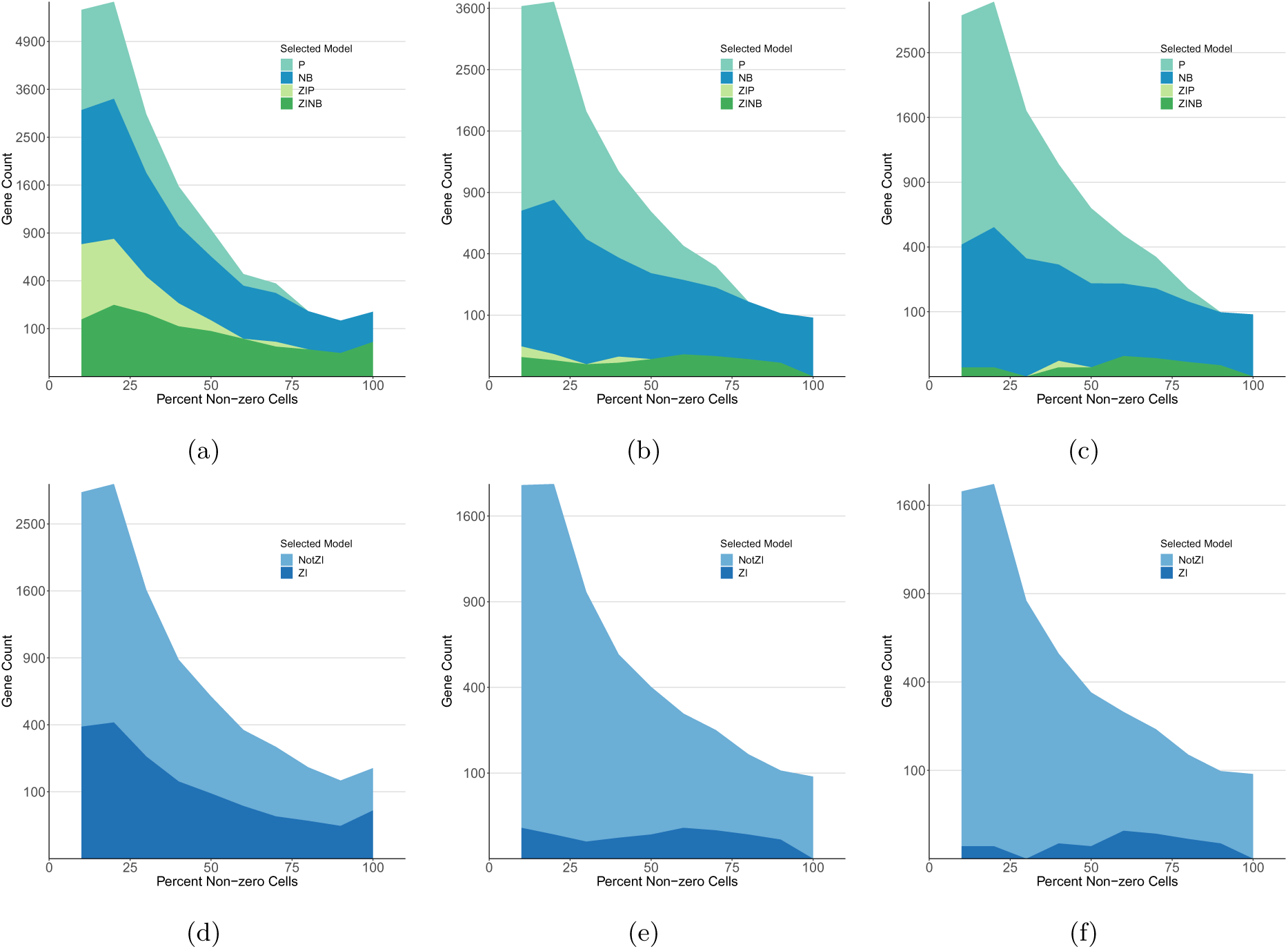
Classification of genes by scRATE model selection applied to heart data. (a) Density plot of model classification of genes across percentages of non-zero cells using scRATE with the 0 SE threshold. (b) As above, for the 2 SE threshold. (c) As above, for the 3 SE threshold. Density plots of scRATE classification collapsed to show only the ZI versus NotZI genes across percentages of non-zero cells with the 0 SE (d), 2 SE (e), and 3 SE (f) thresholds as indicated.

**Supplemental Figure S12:**
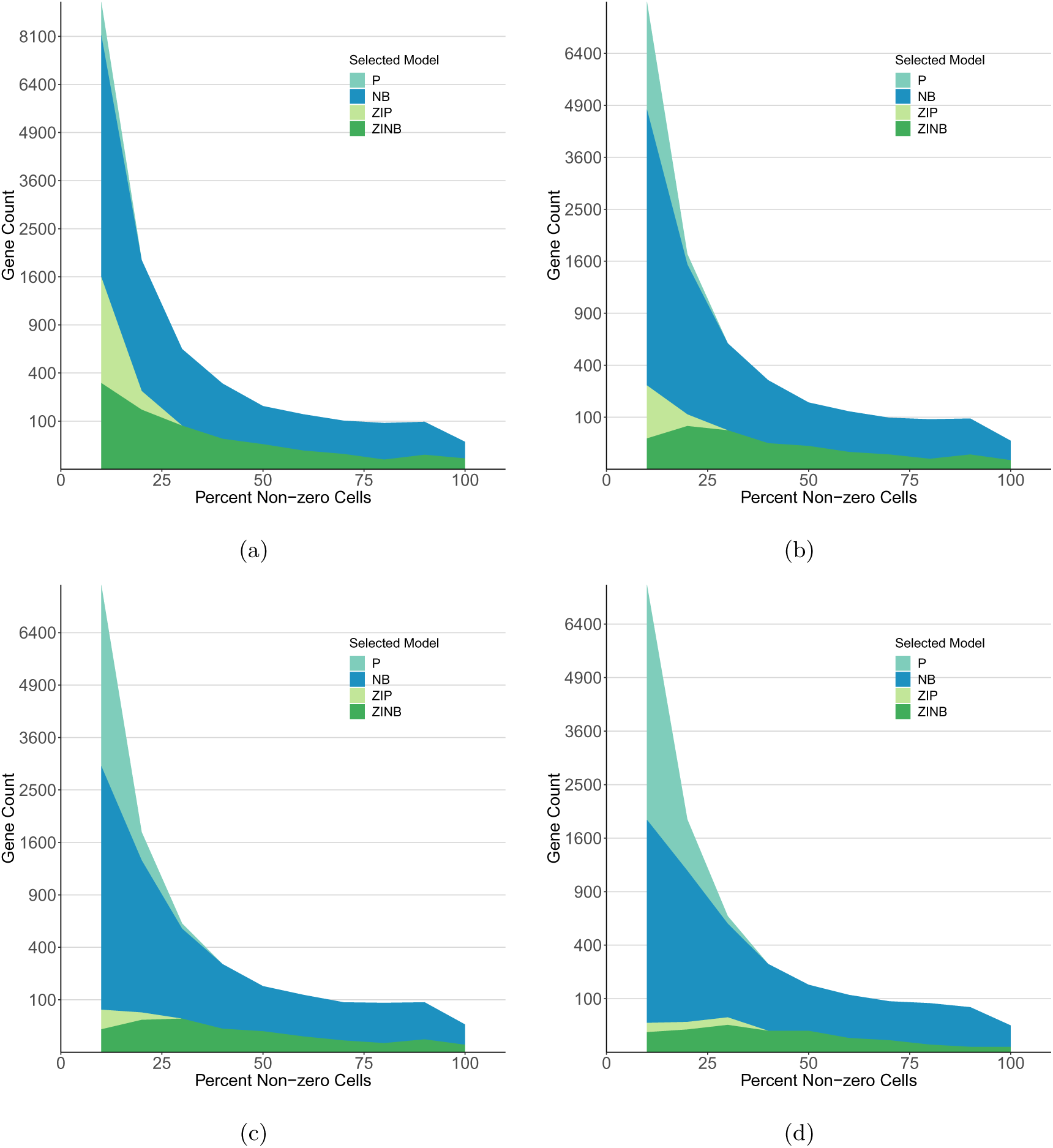
Classification of genes by scRATE model selection applied to the kidney data. (a) Density plot of model classification of genes across percentages of non-zero cells using scRATE with the 0 SE threshold. (b) As above, for the 1 SE threshold, (c) for the 2 SE threshold, and (d) for the 3 SE threshold.

**Supplemental Figure S13:**
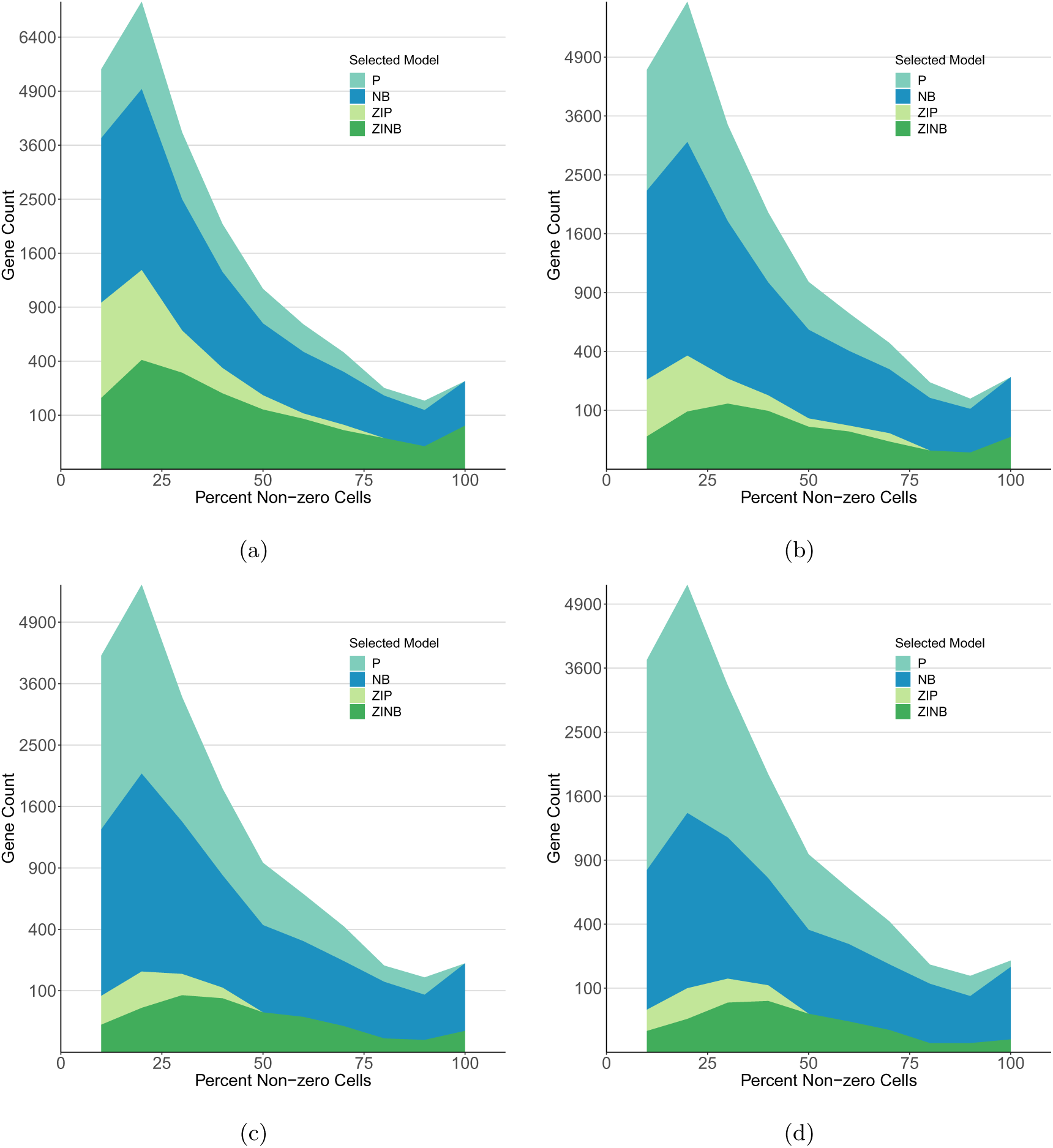
Classification of genes by scRATE model selection applied to the PBMC data. (a) Density plot of model classification of genes across percentages of non-zero cells using scRATE with the 0 SE threshold. (b) As above, for the 1 SE threshold, (c) for the 2 SE threshold, and (d) for the 3 SE threshold.

**Supplemental Figure S14:**
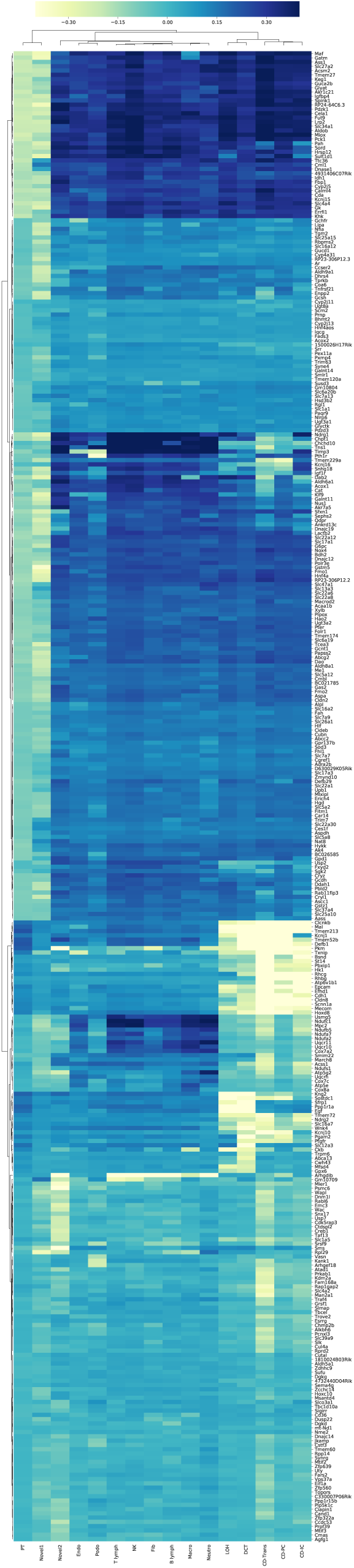
Zeros cluster within specific cell types for ZI genes in the kidney data. A biclustered heatmap of ZI genes (1 SE) by cell types shows the deviation from expected number of cells with zero UMI counts. Dark shading indicates an excess of zeros and light shading indicates that the cell type has fewer cells with zero UMI count than expected.

**Supplemental Figure S15:**
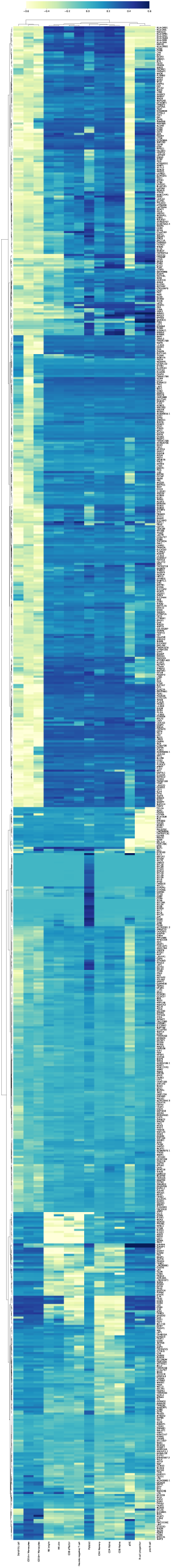
Zeros cluster within specific cell types for ZI genes in the PBMC data. A biclustered heatmap of ZI genes (1 SE) by cell types shows the deviation from expected number of cells with zero UMI counts. Dark shading indicates an excess of zeros and light shading indicates that the cell type has fewer cells with zero UMI count than expected.

**Supplemental Figure S16:**
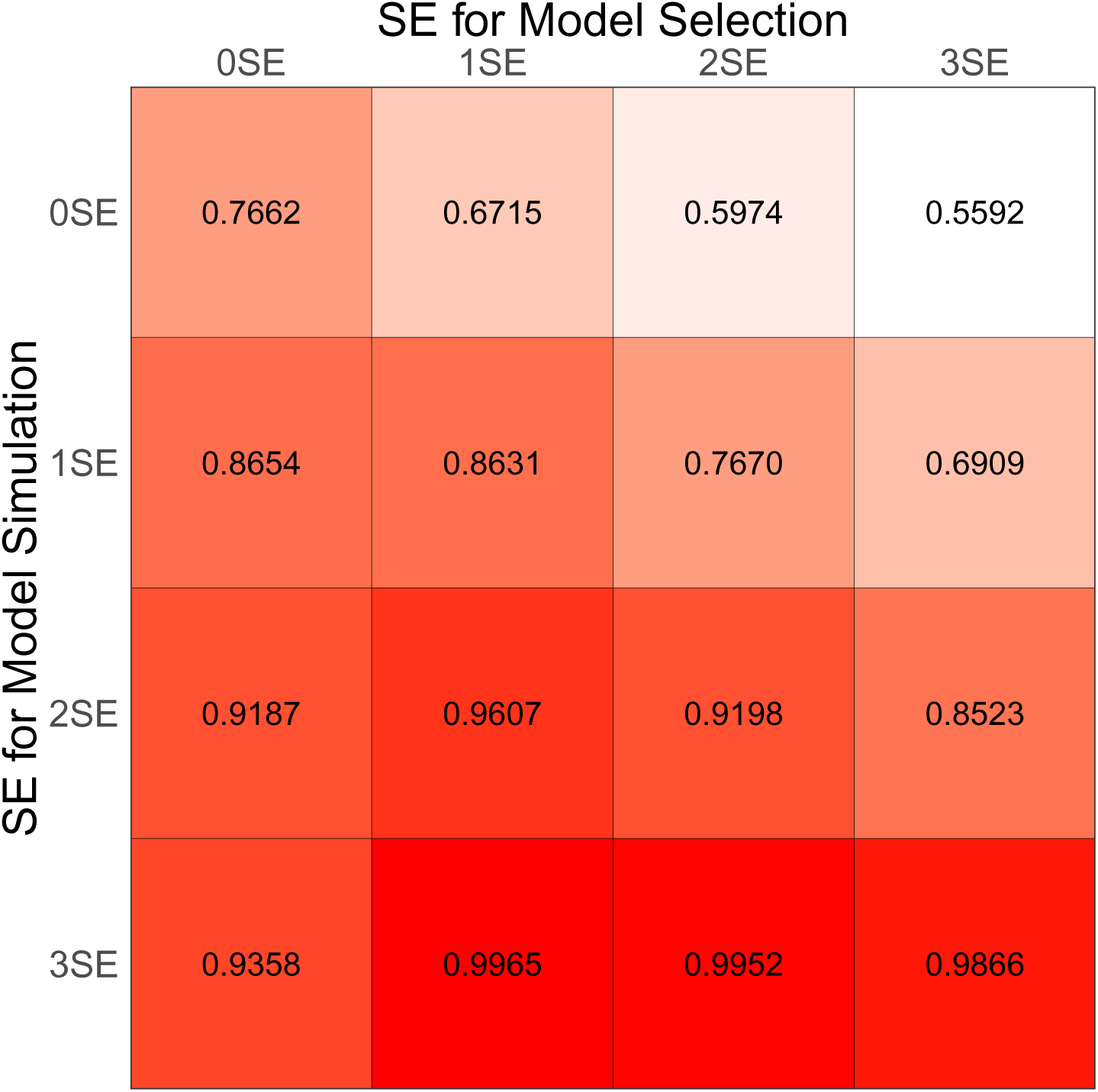
AUC of different model selection thresholds with the simulation based on the kidney data. In order to identify the optimal threshold as in Supplemental Figure S3, we evaluated AUC with simulated gene sets for which distributions are selected from the kidney data with the 0, 1, 2, and 3 SE thresholds (rows). For each simulated gene set, we performed scRATE classification with the 0, 1, 2, and 3 SE thresholds (columns). We find that the 1 SE threshold (the second column) is again robust and performs relatively well across the simulated gene sets.

**Supplemental Figure S17:**
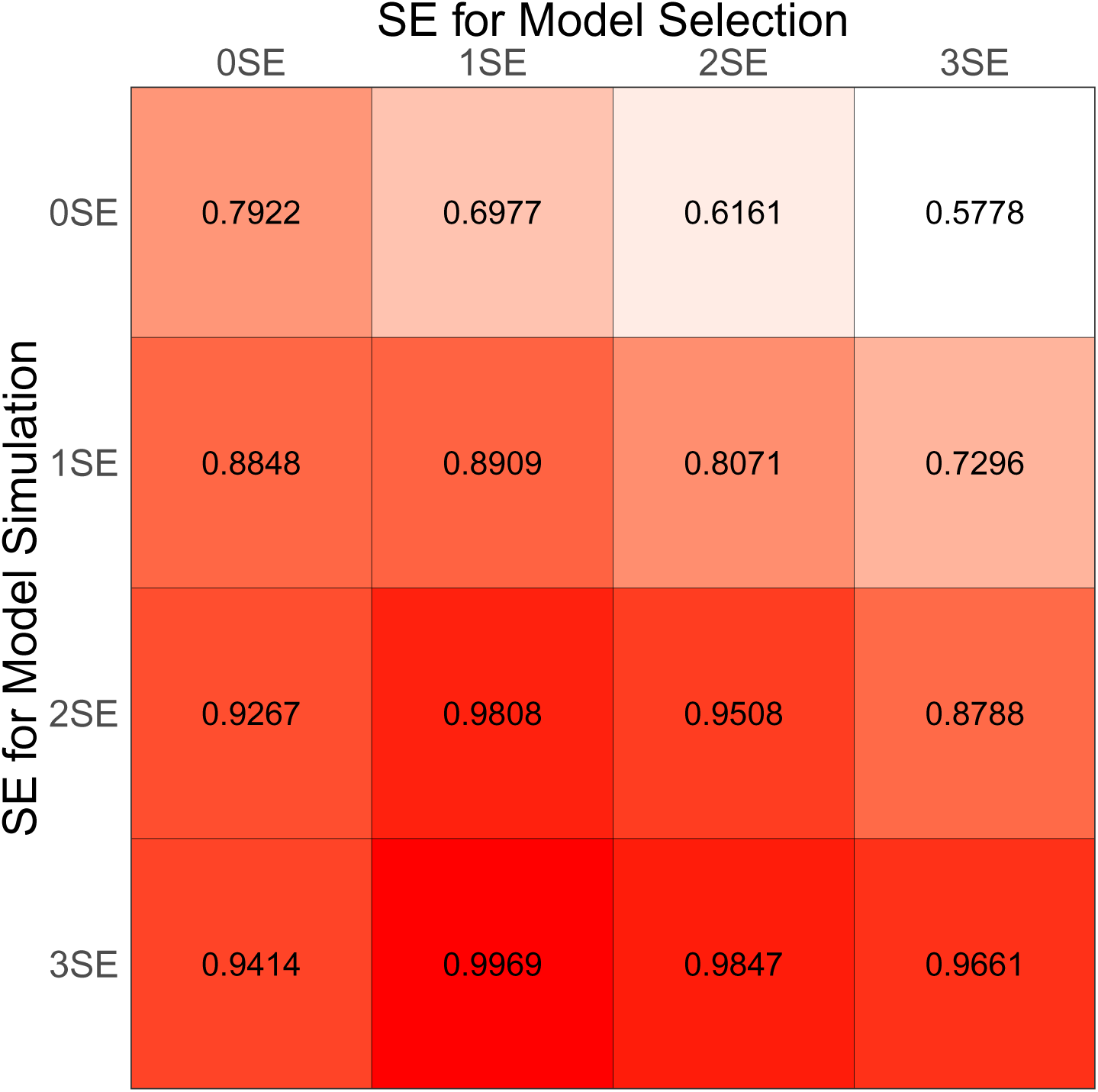
AUC of different model selection thresholds with the simulation based on the PBMC data. In order to identify the optimal threshold as in Supplemental Figure S3 and S16, we evaluated AUC with simulated gene sets for which distributions are selected from the PBMC data with the 0, 1, 2, and 3 SE thresholds (rows). For each simulated gene set, we performed scRATE classification with the 0, 1, 2, and 3 SE thresholds (columns). We find again that the 1 SE threshold (the second column) is robust and performs relatively well across the simulated gene sets.

**Table 2:**
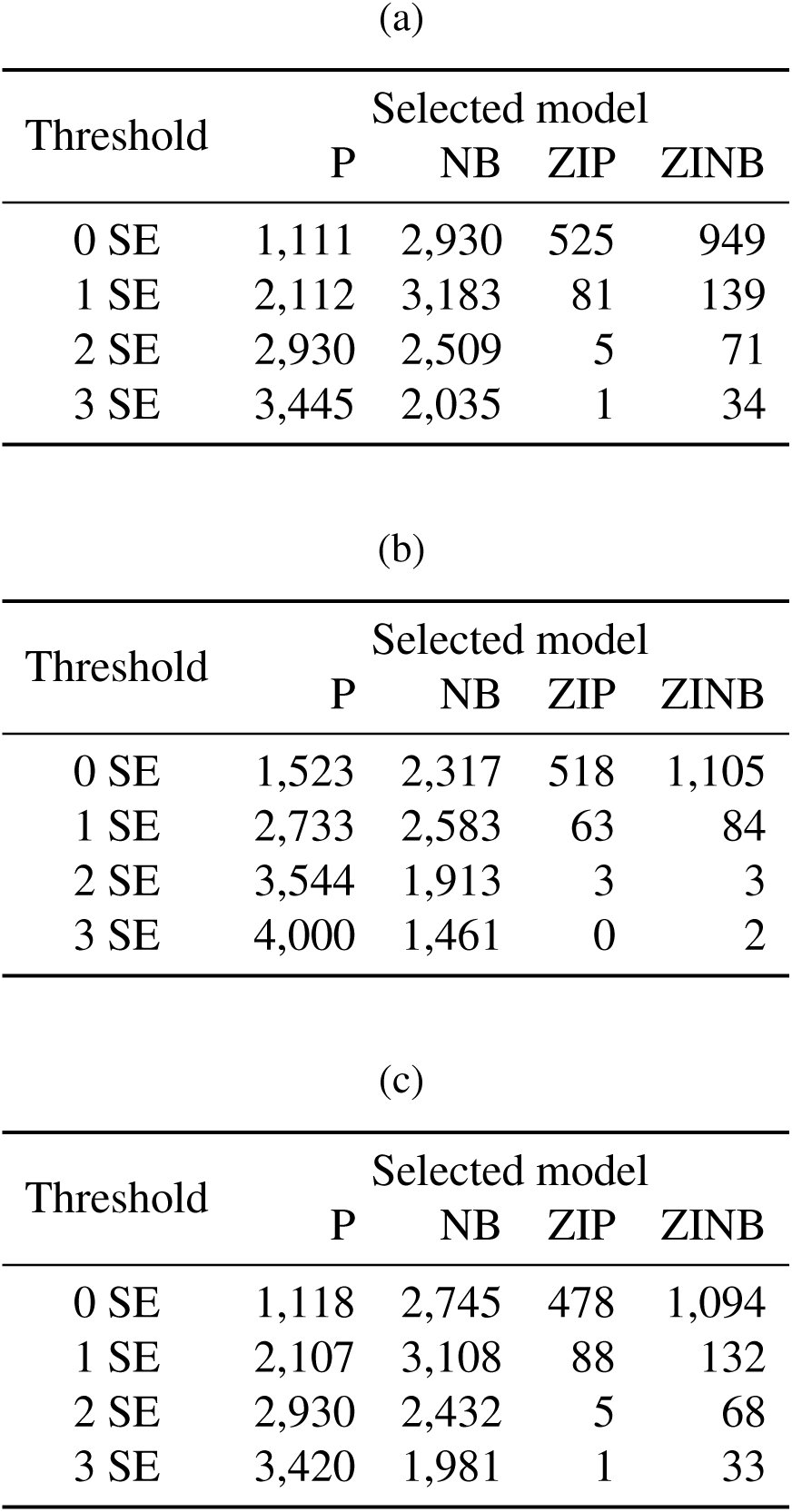
`scRATE` classification of genes in the heart data. Genes were classified as one of four count models (P, NB, ZIP, or ZINB) using four levels of stringency (0 SE, 1 SE, 2 SE, or 3 SE). Table shows number of genes in each category using a GLM with only the offset term to account for cell sequencing depth (a), using a GLM that also includes cell type as an explanatory covariate (b), and using a GLM that includes offset as well as a randomly shuffled cell type as a covariate (c).

| (a) |  |  |  |  |
| --- | --- | --- | --- | --- |
| Threshold | Selected model |  |  |  |
|  | P | NB | ZIP | ZINB |
| 0 SE | 1,111 | 2,930 | 525 | 949 |
| 1 SE | 2,112 | 3,183 | 81 | 139 |
| 2 SE | 2,930 | 2,509 | 5 | 71 |
| 3 SE | 3,445 | 2,035 | 1 | 34 |

| (b) |  |  |  |  |
| --- | --- | --- | --- | --- |
| Threshold | Selected model |  |  |  |
|  | P | NB | ZIP | ZINB |
| 0 SE | 1,523 | 2,317 | 518 | 1,105 |
| 1 SE | 2,733 | 2,583 | 63 | 84 |
| 2 SE | 3,544 | 1,913 | 3 | 3 |
| 3 SE | 4,000 | 1,461 | 0 | 2 |

| (c) |  |  |  |  |
| --- | --- | --- | --- | --- |
| Threshold | Selected model |  |  |  |
|  | P | NB | ZIP | ZINB |
| 0 SE | 1,118 | 2,745 | 478 | 1,094 |
| 1 SE | 2,107 | 3,108 | 88 | 132 |
| 2 SE | 2,930 | 2,432 | 5 | 68 |
| 3 SE | 3,420 | 1,981 | 1 | 33 |

**Table 1:** Error rates and power of `scRATE` classification. Estimated false positive (FP, type I error) and true positive (TP, power) classification rates estimated from simulated data at average depth of 10,000 or 50,000 UMIs per cell. See Methods for details.

| Sequencing<br>depth | Threshold |  |  |
| --- | --- | --- | --- |
|  | 0 SE | 1 SE | 2 SE |
| 10k | 0.219 | 0.017 | 0.002 |
| 50k | 0.231 | 0.034 | 0.001 |

(b)
| Sequencing<br>depth | Threshold |  |  |
| --- | --- | --- | --- |
|  | 0 SE | 1 SE | 2 SE |
| 10k | 0.788 | 0.587 | 0.461 |
| 50k | 0.879 | 0.776 | 0.655 |

**Supplemental Table S1:** Properties of genes classified by scRATE

(a)
| 0 SE model | NumBest | mNumZeros | medNumZeros | mPctNZ | medPctNZ | medAvgExpr | mAvgExpr |
| --- | --- | --- | --- | --- | --- | --- | --- |
| P | 1,111 | 8,366 | 8,710 | 20.4 | 17.1 | 0.192 | 0.251 |
| NB | 2,930 | 7,384 | 8,159 | 29.7 | 22.4 | 0.299 | 0.756 |
| ZIP | 525 | 8,581 | 8,865 | 18.3 | 15.6 | 0.178 | 0.223 |
| ZINB | 949 | 6,467 | 7,293 | 38.5 | 30.6 | 0.415 | 1.27 |

(b)
| 1 SE model | NumBest | mNumZeros | medNumZeros | mPctNZ | medPctNZ | medAvgExpr | mAvgExpr |
| --- | --- | --- | --- | --- | --- | --- | --- |
| P | 2,112 | 8,340 | 8,694 | 20.6 | 17.3 | 0.196 | 0.257 |
| NB | 3,183 | 7,119 | 7,924 | 32.3 | 24.6 | 0.338 | 0.889 |
| ZIP | 81 | 8,216 | 8,465 | 21.8 | 19.4 | 0.229 | 0.287 |
| ZINB | 139 | 4,553 | 4,435 | 56.7 | 57.8 | 1.39 | 3.05 |

(c)
| 2 SE model | NumBest | mNumZeros | medNumZeros | mPctNZ | medPctNZ | medAvgExpr | mAvgExpr |
| --- | --- | --- | --- | --- | --- | --- | --- |
| P | 2,930 | 8,285 | 8,676 | 21.2 | 17.4 | 0.200 | 0.268 |
| NB | 2,509 | 6,733 | 7,612 | 35.9 | 27.6 | 0.412 | 1.16 |
| ZIP | 5 | 8,546 | 9,127 | 18.7 | 13.1 | 0.148 | 0.249 |
| ZINB | 71 | 5,074 | 4,455 | 51.7 | 57.6 | 1.41 | 1.89 |

**Supplemental Table S2:** Mean square errors of mean and overdispersion parameters for the genes in the heart data (a) NB simulation before cell type adjustment, (b) ZINB simulation before cell type adjustment, (c) NB simulation using cell type as covariate, and (d) ZINB simulation using cell type as covariate.

| (a) |  |  | (b) |  |  |
| --- | --- | --- | --- | --- | --- |
| NB simulation | NB | ZINB | ZINB simulation | NB | ZINB |
| mean | 0.00153 | 0.00811 | mean | 0.01385 | 0.00587 |
| overdispersion | 0.0166 | 0.2817 | overdispersion | 0.24941 | 0.14867 |

| (c) |  |  | (d) |  |  |
| --- | --- | --- | --- | --- | --- |
| NB simulation | NB | ZINB | ZINB simulation | NB | ZINB |
| mean | 0.00548 | 0.00777 | mean | 0.00812 | 0.00705 |
| overdispersion | 0.009 | 0.052 | overdispersion | 0.0444 | 0.0165 |

**Supplemental Table S3:** Number of models called for the genes in the kidney data without using cell type as covariate (a), using cell type as covariate (b), and using cell type as covariate after randomly shuffling cell type labels (c).

| Threshold | Selected model |  |  |  |
| --- | --- | --- | --- | --- |
|  | P | NB | ZIP | ZINB |
| 0 SE | 48 | 3,946 | 500 | 666 |
| 1 SE | 434 | 4,378 | 109 | 239 |
| 2 SE | 1,225 | 3,760 | 16 | 159 |
| 3 SE | 2,024 | 3,023 | 7 | 106 |

(b)
| Threshold | Selected model |  |  |  |
| --- | --- | --- | --- | --- |
|  | P | NB | ZIP | ZINB |
| 0 SE | 104 | 3,654 | 485 | 917 |
| 1 SE | 730 | 4,221 | 117 | 92 |
| 2 SE | 1,807 | 3,311 | 9 | 33 |
| 3 SE | 2,679 | 2,471 | 0 | 10 |

(c)
| Threshold | Selected model |  |  |  |
| --- | --- | --- | --- | --- |
|  | P | NB | ZIP | ZINB |
| 0 SE | 52 | 3,820 | 459 | 829 |
| 1 SE | 447 | 4,367 | 111 | 235 |
| 2 SE | 1,272 | 3,711 | 18 | 159 |
| 3 SE | 2,053 | 2,995 | 7 | 105 |

**Supplemental Table S4:**
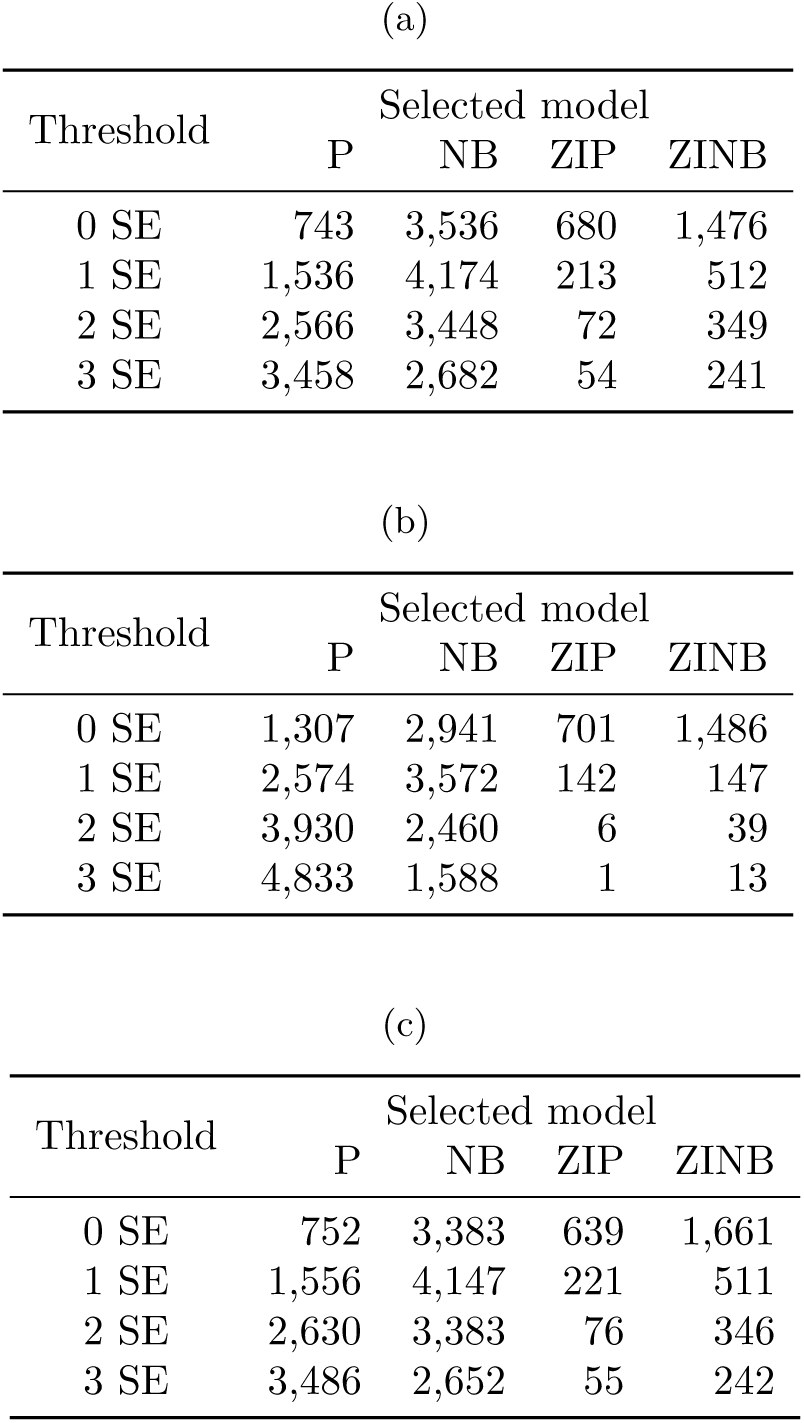
The number of models called for the genes in the PBMC data without using cell type as covariate (a), using cell type as covariate (b), and using cell type as covariate after randomly shuffling cell type labels (c).

## References

1. [10X Genomics, 2018]10X Genomics, 2018. 10k PBMCs from a Healthy Donor (v3 chemistry). https://support.10xgenomics.com/single-cell-gene-expression/datasets/3.0.0/pbmc_10k_v3. Accessed: Oct. 13th, 2019.

2. [Andrews and Hemberg, 2019]Andrews, T. and Hemberg, M., 2019. False signals induced by single-cell imputation [version 2; peer review: 4 approved]. F1000Research, 7(1740).

3. [Butler et al., 2018]Butler, A., Hoffman, P., Smibert, P., Papalexi, E., and Satija, R., 2018. Integrating single-cell transcriptomic data across different conditions, technologies, and species. Nature Biotechnology, 36(5):411–420.

4. [Bürkner, 2017]Bürkner, P.-C., 2017. brms: An R package for Bayesian multilevel models using Stan. Journal of Statistical Software, 80(1):1–28.

5. [Bürkner, 2018]Bürkner, P.-C., 2018. Advanced Bayesian multilevel modeling with the R package brms. The R Journal, 10(1):395–411.

6. [Campbell, 2019]Campbell, H., 2019. The consequences of checking for zero-inflation and overdispersion in the analysis of count data.

7. [Cao et al., 2017]Cao, J., Packer, J. S., Ramani, V., Cusanovich, D. A., Huynh, C., Daza, R., Qiu, X., Lee, C., Furlan, S. N., Steemers, F. J., et al., 2017. Comprehensive single-cell transcriptional profiling of a multicellular organism. Science, 357(6352):661–667.

8. [Gelman and Loken, 2014]Gelman, A. and Loken, E., 2014. The statistical crisis in science. American Scientist, 102(6):460–465.

9. [Gong et al., 2018]Gong, W., Kwak, I.-Y., Pota, P., Koyano-Nakagawa, N., and Garry, D. J., 2018. Drimpute: imputing dropout events in single cell rna sequencing data. BMC Bioinformatics, 19(1):220.

10. [Goodrich et al., 2019]Goodrich, B., Gabry, J., Ali, I., and Brilleman, S., 2019. rstanarm: Bayesian applied regression modeling via Stan. R package version 2.19.2.

11. [Goodrich et al., 2020]Goodrich, B., Gabry, J., Ali, I., and Brilleman, S., 2020. rstanarm: Bayesian applied regression modeling via Stan. R package version 2.19.3.

12. [Hicks et al., 2017]Hicks, S. C., Townes, F. W., Teng, M., and Irizarry, R. A., 2017. Missing data and technical variability in single-cell RNA-sequencing experiments. Biostatistics, 19(4):562–578.

13. [Hooten and Hefley, 2019]Hooten, M. B. and Hefley, T. J., 2019. Bringing bayesian models to life. CRC Press, Taylor et Francis.

14. [Islam et al., 2014]Islam, S., Zeisel, A., Joost, S., La Manno, G., Zajac, P., Kasper, M., Lön-nerberg, P., and Linnarsson, S., 2014. Quantitative single-cell rna-seq with unique molecular identifiers. Nature Methods, 11(2):163–166.

15. [Kharchenko et al., 2014]Kharchenko, P. V., Silberstein, L., and Scadden, D. T., 2014. Bayesian approach to single-cell differential expression analysis. Nature Methods, 11(7):740–742.

16. [Klein et al., 2015]Klein, A. M., Mazutis, L., Akartuna, I., Tallapragada, N., Veres, A., Li, V., Peshkin, L., Weitz, D. A., and Kirschner, M. W., 2015. Droplet barcoding for single-cell transcriptomics applied to embryonic stem cells. Cell, 161(5):1187 – 1201.

17. [Li and Li, 2018]Li, W. V. and Li, J. J., 2018. An accurate and robust imputation method scimpute for single-cell rna-seq data. Nature Communications, 9(1):997.

18. [Macosko et al., 2015]Macosko, E. Z., Basu, A., Satija, R., Nemesh, J., Shekhar, K., Gold-man, M., Tirosh, I., Bialas, A. R., Kamitaki, N., Martersteck, E. M., et al., 2015. Highly parallel genome-wide expression profiling of individual cells using nanoliter droplets. Cell, 161(5):1202–1214.

19. [Park et al., 2018]Park, J., Shrestha, R., Qiu, C., Kondo, A., Huang, S., Werth, M., Li, M., Barasch, J., and Suszták, K., 2018. Single-cell transcriptomics of the mouse kidney reveals potential cellular targets of kidney disease. Science, 360(6390):758–763.

20. [Rosenberg et al., 2018]Rosenberg, A. B., Roco, C. M., Muscat, R. A., Kuchina, A., Sample, P., Yao, Z., Graybuck, L. T., Peeler, D. J., Mukherjee, S., Chen, W., et al., 2018. Single-cell profiling of the developing mouse brain and spinal cord with split-pool barcoding. Science, 360(6385):176–182.

21. [Skelly et al., 2018]Skelly, D. A., Squiers, G. T., McLellan, M. A., Bolisetty, M. T., Robson, P., Rosenthal, N. A., and Pinto, A. R., 2018. Single-cell transcriptional profiling reveals cellular diversity and intercommunication in the mouse heart. Cell Reports, 22(3):600 – 610.

22. [Stanley et al., 2019]Stanley, G., Gokce, O., Malenka, R. C., Südhof, T. C., and Quake, S. R., 2019. Discrete and continuous cell identities of the adult murine striatum. bioRxiv,.

23. [Svensson, 2020]Svensson, V., 2020. Droplet scrna-seq is not zero-inflated. Nature Biotechnology, 38(2):147–150.

24. [Townes et al., 2019]Townes, F. W., Hicks, S. C., Aryee, M. J., and Irizarry, R. A., 2019. Feature selection and dimension reduction for single-cell rna-seq based on a multinomial model. Genome Biology, 20(1):295.

25. [Vehtari et al., 2019]Vehtari, A., Gabry, J., Magnusson, M., Yao, Y., and Gelman, A., 2019. loo: Efficient leave-one-out cross-validation and waic for bayesian models. R package version 2.2.0.

26. [Vehtari et al., 2017]Vehtari, A., Gelman, A., and Gabry, J., 2017. Practical bayesian model evaluation using leave-one-out cross-validation and waic. Statistics and Computing, 27:1413–1432.

27. [Zappia and Oshlack, 2018]Zappia, L. and Oshlack, A., 2018. Clustering trees: a visualization for evaluating clusterings at multiple resolutions. GigaScience, 7(7). giy083.

28. [Zeileis et al., 2008]Zeileis, A., Kleiber, C., and Jackman, S., 2008. Regression models for count data in r. Journal of Statistical Software, Articles, 27(8):1–25.

## References

30. [10X Genomics, 2018]10X Genomics, 2018. 10k PBMCs from a Healthy Donor (v3 chemistry). https://support.10xgenomics.com/single-cell-gene-expression/datasets/3.0.0/pbmc_10k_v3. Accessed: Oct. 13th, 2019.

31. [Park et al., 2018]Park, J., Shrestha, R., Qiu, C., Kondo, A., Huang, S., Werth, M., Li, M., Barasch, J., and Susztáak, K., 2018. Single-cell transcriptomics of the mouse kidney reveals potential cellular targets of kidney disease. Science, 360(6390):758–763.

32. [Stuart et al., 2019]Stuart, T., Butler, A., Hoffman, P., Hafemeister, C., Papalexi, E., Mauck, W. M., Hao, Y., Stoeckius, M., Smibert, P., and Satija, R., et al., 2019. Comprehensive integration of single-cell data. Cell, 177(7):1888 –1902.e21.

